# Platform-Agnostic CellNet (PACNet) enables cross-study meta-analysis of cell fate engineering protocols

**DOI:** 10.1101/2022.09.07.506886

**Authors:** Emily K.W. Lo, Jeremy Velazquez, Da Peng, Chulan Kwon, Mo R. Ebrahimkhani, Patrick Cahan

**Affiliations:** Department of Biomedical Engineering, Johns Hopkins University, Baltimore, MD, 21205, United States; Institute for Cell Engineering, Johns Hopkins University, Baltimore, MD, 21205, United States; Department of Pathology, School of Medicine, University of Pittsburgh, Pittsburgh, 15213, PA, USA; Pittsburgh Liver Research Center, University of Pittsburgh, Pittsburgh, 15261, PA, USA; Department of Medicine, Johns Hopkins University, Baltimore, MD, 21205, United States; Department of Bioengineering, Swanson School of Engineering, University of Pittsburgh, Pittsburgh, 15261, PA, USA; McGowan Institute for Regenerative Medicine, University of Pittsburgh, Pittsburgh, 15219, PA, USA

**Keywords:** Cell Engineering, Directed Differentiation, Transdifferentiation, Organoids, Cardiomyocytes, Hepatocytes, Heart, Liver, Computational Biology, Transcriptomics

## Abstract

The optimization of cell fate engineering protocols requires evaluating their fidelity, efficiency, or both. We previously adopted CellNet, a computational tool to quantitatively assess the transcriptional fidelity of engineered cells and tissues as compared to their in vivo counterparts based on bulk RNA-Seq. However, this platform and other similar approaches are sensitive to experimental and analytical aspects of transcriptomics methodologies. This makes it challenging to capitalizing on the expansive, publicly available sets of transcriptomic data that reflect the diversity of cell fate engineering protocols. Here, we present Platform-Agnostic CellNet (PACNet), which extends the functionality of CellNet by enabling the assessment of transcriptional profiles in a platform-agnostic manner, and by enabling the comparison of user-supplied data to panels of engineered cell types from state-of-the-art protocols. To demonstrate the utility of PACNet, we evaluated a range of cell fate engineering protocols for cardiomyocytes and hepatocytes. Through this analysis, we identified the best-performing methods, characterized the extent of intra-protocol and inter-lab variation, and identified common off-target signatures, including a surprising neural and neuroendocrine signature in primary liver-derived organoids. Finally, we made our tool accessible as a user-friendly web application that allows users to upload their own transcriptional profiles and assess their protocols relative to our database of reference engineered samples.

**Highlights:**

• The development of Platform-Agnostic CellNet (PACNet) that classifies engineered cell populations from transcriptome data regardless of profiling method or transcript abundance estimation method
• PACNet enables cross-study comparisons of cell fate engineering protocols
• Comparison of cardiomyocyte engineering protocols emphasizes metabolic selection as a key step in achieving a strong cardiomyocyte fate.
• PACNet identifies an unexpected off-target neural and neuroendocrine signature in primary liver-derived organoids.

**eTOC Blurb:** Cahan and colleagues created a computational resource, PACNet, which evaluates the fidelity of cell engineering expression profiles in a platform-agnostic manner to facilitate cross-protocol benchmarking. Examining state-of-the-field cardiomyocyte and hepatocyte derivation protocols, they identified that two techniques in cardiomyocyte engineering best increase cardiac identity and that an off-target neural/neuroendocrine signature in primary liver-derived organoids may reflect a cholangiopathic signature.

**Graphical abstract:** 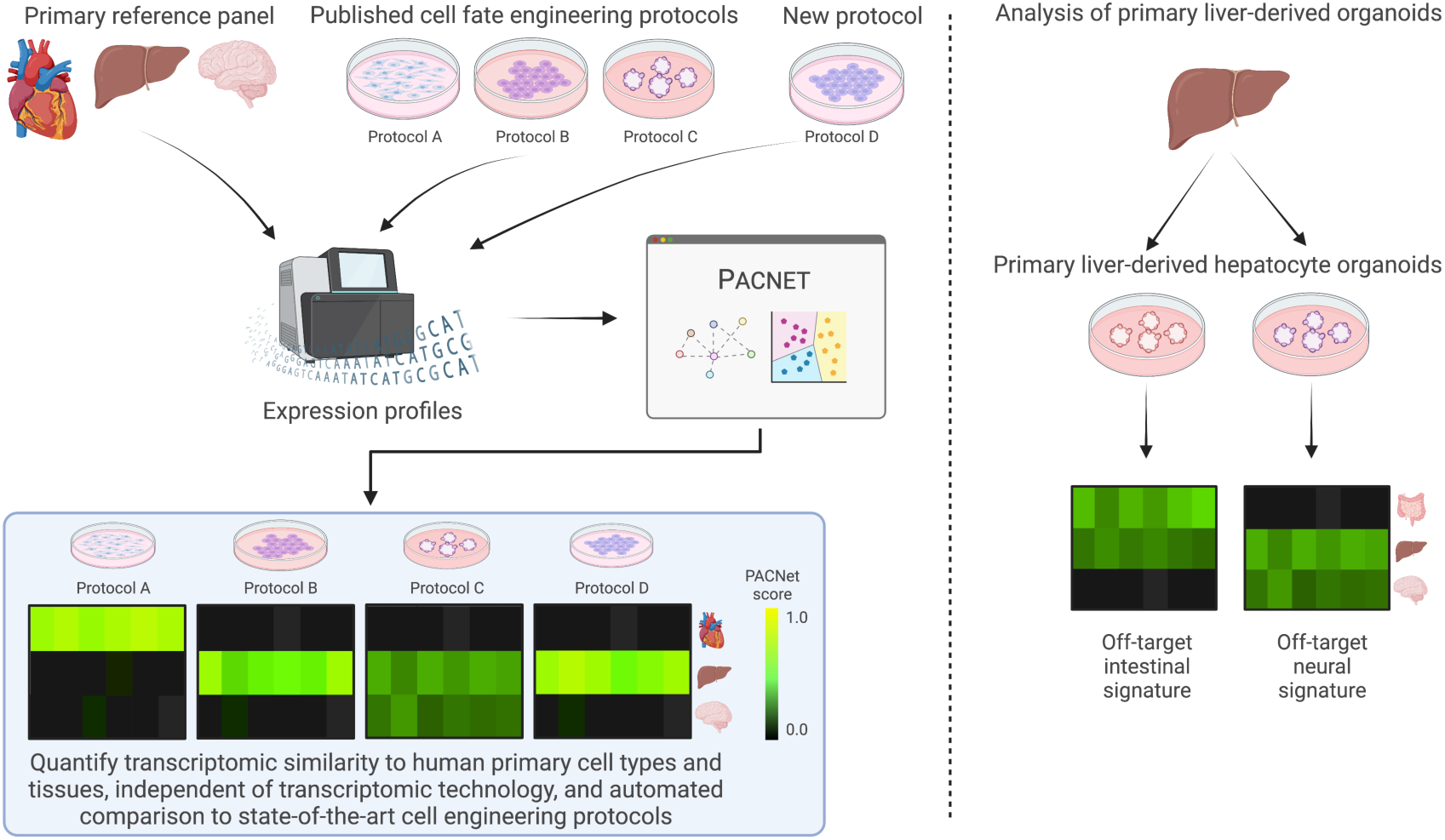

## Introduction

Key milestones in advancing the field of cell fate engineering (CFE) include the discovery of direct conversion (Davis et al.), the derivation of mouse and human embryonic stem cells (ESCs) (Evans and Kaufman, 1981; Martin, 1981; Thomson et al., 1998), and the innovation of induced pluripotent stem cells (iPSCs) (Takahashi and Yamanaka, 2006; Takahashi et al., 2007). Collectively, these advancements have enabled the development of protocols to derive numerous cell and tissue types. CFE is used in a range of applications, from regenerative medicine, to disease modeling, to drug discovery and screening (Robinton and Daley, 2012). Due to the potential import of these applications, the number of investigators and studies generating, optimizing, and applying CFE methods has multiplied rapidly.

Optimizing cell and tissue engineering methods implicitly necessitates the evaluation of protocol performance. This is often done empirically by verifying the expression of canonical markers at both the protein and RNA level, for instance alpha-1 antitrypsin and hepatocyte nuclear factors in hepatocytes (Ma et al., 2013) and cardiac troponin in cardiomyocytes (Ieda et al., 2010), or through in vitro morphological and functional assays, for example measuring glycogen storage and albumin secretion in hepatocytes (Ma et al., 2013), calcium flux and spontaneous beating in cardiomyocytes (Ieda et al., 2010), and sodium currents and spontaneous postsynaptic currents in neurons (Kang et al., 2017). The ideal empirical assay for evaluating cell type identity is transplantation into and rescue of an absent or ablated *in vivo* function. The most notable functional assays historically include the teratoma (Heins et al., 2004; Thomson et al., 1998) and tetraploid complementation (Kang et al., 2009; Tam and Rossant, 2003) assays for assessing pluripotency, as well as the engraftment of engineered hematopoietic stem cells to repopulate the bone marrow niche (Vo and Daley, 2015). Approaches that rely on genome-wide profiling have complemented these functional assays as they are often easier and faster to execute, and they can identify aberrant signatures. Specialized computational tools for evaluating the fidelity of CFE protocols have been developed, for example TeratoScore, PluriTest, and ScoreCard for assessing pluripotency (Allison et al., 2018; Avior et al., 2015a; Tsankov et al., 2015) and KeyGenes for assessing developmental stage using a fetal tissue atlas (Roost et al., 2015). We also previously developed one such tool, CellNet (Cahan et al., 2014; Radley et al., 2017), which compares transcriptional profiles of author-derived engineered cells to those of primary adult samples in order to assess the performance of CFE protocols. These methods all compare query sample profiles to the primary *in vivo* profiles on which they are trained.

These current computational approaches to evaluate the products of CFE are limited in that they are typically optimized for and deployed in a study-specific manner. Furthermore, while single-cell RNA-Seq comparisons of engineered versus *in vivo* cells can distinguish hybrid identity from population heterogeneity (Han et al., 2020; Shapiro et al., 2013; Tang et al., 2011), the methods typically used to integrate scRNAseq data from multiple sources (i.e. when comparing CFE products to a previously published single cell atlas) tend to over-merge cells with distinct cell states (Babcock et al 2021) (Richards et al., 2021). These issues make it challenging to fairly compare CFE products across protocols and studies. Such cross-study comparisons of CFE would help to determine the extent to which new CFE protocols improve fidelity as compared to standard methods in the field, and to investigate intra-protocol variability.

To address this deficiency, we developed a platform-agnostic CellNet (PACNet), which provides cross-study comparisons examining bulk transcriptomic profiles of investigator-engineered cells derived from state-of-the-art protocols. PACNet is agnostic to the platform of expression profiling, including microarray and Illumina- and ION torrent-based RNA-Seq, as well as the method of preprocessing raw sequencing data into gene expression estimates. We have compiled and evaluated a database of publicly available bulk human gene expression data from 104 CFE experiments, totaling more than 2100 samples across seven cell/tissue types. Using this platform, we quantitatively evaluate the most common, most consistent, and best-performing protocols for two of the most prolific engineered tissue types: heart and liver. We identify common off-target signatures across heart and liver engineering protocols and reveal an unexpected neural and neuroendocrine signature in primary liver-derived organoids. Finally, we have created a user-friendly web application (http://cahanlab.org/resources/agnosticCellNet_web/) through which investigators can upload gene expression data to compare their engineered cell types to our entire database of engineered reference samples. Visitors to the site can also explore the results of PACNet as applied to all 104 CFE data sets. Users also have the option of downloading our database of reference samples and running our code locally (https://github.com/pcahan1/CellNet). This tool will serve as a valuable resource for investigators to better evaluate the efficacy and performance of CFE protocols within the context of the field.

## Results

### Platform-agnostic CellNet classifiers are precise and sensitive

To train a platform-agnostic CellNet, we first mined NCBI Gene Expression Omnibus (GEO) for bulk RNA-Seq profiles of primary, healthy human tissue samples from 14 cell types: B cells, endothelial cells, ESCs, fibroblasts, heart, hematopoietic stem and progenitor cells (HSPCs), intestine/colon, kidney, liver, lung, monocytes/macrophages, brain, skeletal muscle, and T cells. This process resulted in more than 1400 samples across the 14 cell types (Table S1). PACNet trained on these healthy human samples performed well when applied to a subset of held-out samples, with an average classification score (corresponding to the labeled cell type) of 0.965 among all held-out samples, with cell type-specific averages ranging from 0.872 (kidney) to 0.996 (neuron) (Fig. 1A). We also evaluated classifier performance via precision-recall (PR) curves, summarized as area under the PR curve (AUPR). The AUPR among cell types averaged 0.9981 and ranged from 0.9962 (kidney) to 0.9996 (monocyte/macrophage) (Fig S1A-B).

**Figure 1.**
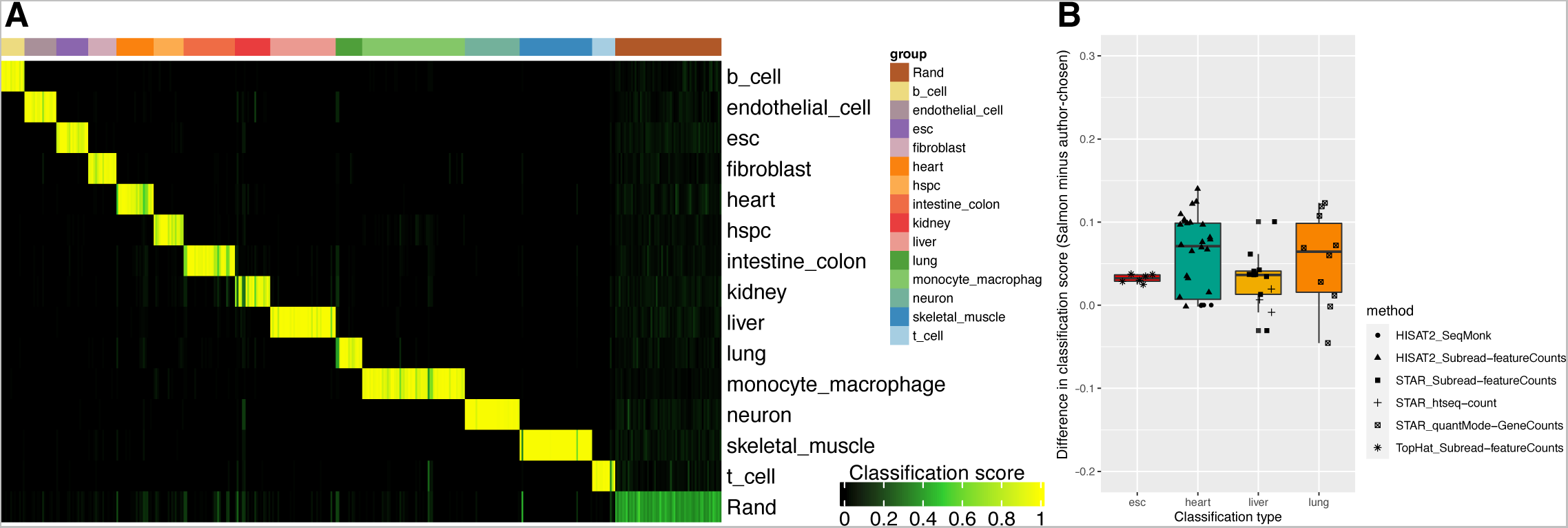
Platform-Agnostic CellNet classifier and preprocessing performance. See also Figure S1. (A) Heatmap of classification scores of held-out samples for classifier validation. Each column represents a held-out bulk RNA-seq sample; each row represents a cell/tissue type-specific top pairs random forest classifier. Classification score of 1 indicates 100% probability that a given sample is indistinguishable from its in vivo training counterparts. Rand: random. (B) Difference in classification scores based on in-house vs author-performed RNA-seq alignment and quantification pipelines for cell fate engineering studies. Only *in vivo* positive controls from iPSC, heart, liver, and lung cell fate engineering studies were analyzed. Method legend: tools used for author-performed RNA-seq alignment and quantification. Mean difference of 0.0495 across all comparisons; no difference in scores was greater than 0.14.

To create a reference database broadly representative of major cell fate protocols, we again mined GEO for publicly available bulk gene expression data. We aimed to identify a variety of protocol types within each cell or tissue type, including directed differentiation, transdifferentiation, and organoid derivation protocols. We also sought to compile studies using a variety of expression profiling platforms, including microarray, both Illumina- and ION torrent-based RNA-Seq, GRO-seq, and TRAP-seq. Altogether we gathered 104 CFE experiments for heart (24 studies), hematopoietic stem and progenitor cells (5), intestine/colon (12), liver (25), lung (5), neuron (21), and skeletal muscle (9), totaling more than 2100 samples (Table S2).

As our goal was to make a metric of CFE that would be well-calibrated across studies that use different preprocessing methods, we compared the performance of PACNet on samples preprocessed using our RNA-Seq quantification method of choice, the alignment-free tool Salmon (Patro et al., 2017), to those using various other author-performed alignment and quantification preprocessing methods, including HISAT2 (Kim et al., 2019), STAR (Dobin et al., 2013), TopHat (Trapnell et al., 2009), Subread’s featureCounts (Liao et al., 2014), and HTSeq’s htseq-count (Anders et al., 2015). We preprocessed each of the held-out primary ESC, heart, liver, and lung samples with Salmon and compared the resulting classification scores against those resulting from the corresponding author-chosen method. There was little difference between classification scores, with a mean difference of 0.0495 across all four evaluated cell types, and no differences greater than 0.14. (Fig. 1B, Table S3). Previous benchmarking studies have shown that although alignment-based methods have an advantage in detecting small RNAs (Wu et al., 2018a), alignment-free methods such as Salmon and Kallisto have a slightly higher concordance with ground truth expression (based on known abundance of exogenous spike-ins and simulated data) and are more computationally efficient than alignment-based methods (Bray et al., 2016; Everaert et al., 2017; Yi et al., 2018). We attribute the robustness of PACNet cell fate assessment – against variations in profiling platform, preprocessing (alignment and quantification) pipeline, and even counts vs per million quantification – to the previously described top-scoring pairs algorithm used in our PACNet classifier (Geman et al., 2004; Peng et al., 2020; Tan and Cahan, 2019), as it compares the relative expression of gene pairs within samples rather than absolute or normalized expression among samples.

To allow the use of different preprocessing methods in user-specific studies, we had to handle the situation where datasets do not share all of the same genes. To achieve this, we trained one PACNet classifier per CFE target cell/tissue type using genes common to both the reference data and all of the data of the studies relevant for that target cell/tissue type. (NB: When new query data is analyzed, our web platform will automatically train a new classifier if genes in the existing classifier are not in the query data and will re-analyze existing CFE studies to enable fair comparison to the new CFE data.) We again used AUPR to evaluate the performance of classifiers trained only on overlapping genes and queried with the same held-out subset of primary healthy human samples. The number of overlapping genes ranged from 5,909 among engineered intestine/colon studies to 20,600 among engineered skeletal muscle studies. Across all classifiers (excluding the ‘random’ category), the mean AUPR remained at 0.9975 compared to the 0.9981 AUPR achieved with a full set of 34,934 genes. The AUPR across all classifiers ranged from 0.9763 (kidney classifier trained with 15,288 common genes among liver studies) to 0.9997 (monocyte/macrophage classifier trained with 12,827 common genes among lung studies) (Fig. S1B), demonstrating the robustness of the top-pairs classifier against variations in available genes.

Cardiomyocytes and hepatocytes are frequently the target of CFE efforts because of their potential applications in toxicity screening and in regenerative medicine (Buikema et al., 2013; Jin et al., 2021; Karakikes et al., 2015; Laflamme and Murry, 2011; Li et al., 2021; Schwartz et al., 2014). In the following sections, we used PACNet to identify the quantify the transcriptional fidelity of common CFE methods, to quantify the extent and frequency of off-target effects, and to explore the biological pathways that distinguish CFE protocols.

### Engineered cardiomyocyte panel emphasizes metabolic selection as a key step

We first identified common derivation protocols among cardiomyocyte (CM) CFE studies (Fig. 2A). Two commonly used CM directed differentiation monolayer protocols, which we hereafter denote by the first authors: Burridge (Burridge et al., 2014, 2015) and Lian (Lian et al., 2012, 2013), begin with GSK3 inhibition with CHIR99021 for 48-72 hours to induce mesoderm development, followed by Wnt inhibition for 48 hours using small molecule PORCN inhibitors (Burridge uses Wnt-C59, Lian uses IWP-2), to generate cardiac mesoderm. Subsequently the cultures undergo standard media changes with either RPMI + L-ascorbic acid 2-phosphate + recombinant human albumin (Burridge), or RPMI + B-27 (Lian). Beating or contractile cells are observed at approximately day 7. At day 10 in the Burridge protocol CMs are subjected to metabolic selection via glucose deprivation and/or sodium DL-lactate supplementation, based on a finding by Tohyama and colleagues (Tohyama et al., 2013), which purifies up to 95% TNNT2+ (troponin T-positive) cells. Differentiated, contractile CMs generated from both protocols can be maintained in culture for more than 6 months. Several studies append the metabolic selection step from Burridge to the Lian protocol (Ang et al., 2016; Banovich et al., 2018; Cyganek et al., 2018; Hookway et al., 2019; Jaffré Fabrice et al., 2019; Strober et al., 2019); we designate these as “Combined”. Another monolayer protocol is remarkably simple, comprising a one-step, 24hr treatment with activin, followed by at least 8 days of culture in just RPMI + B-27 (Estarás et al., 2017; Hsu et al., 2018). We denote these studies as “Activin-based.” Alternative protocols by Yang et al (Yang et al., 2008) and Lee et al (Lee et al., 2017), which we denote as “Embryoid Body” (EB) protocols, use recombinant growth factors in lieu of small molecule inhibitors for differentiation, beginning with the formation of EBs using BMP4, followed by primitive streak induction using BMP4, bFGF, and Activin A, cardiac mesoderm induction using VEGF and Wnt-inhibiting factor DKK1, and finally CM specification using VEGF, DKK1, and bFGF. Finally, we denote commercially purchased CMs as “iCell” CMs.

**Figure 2.**
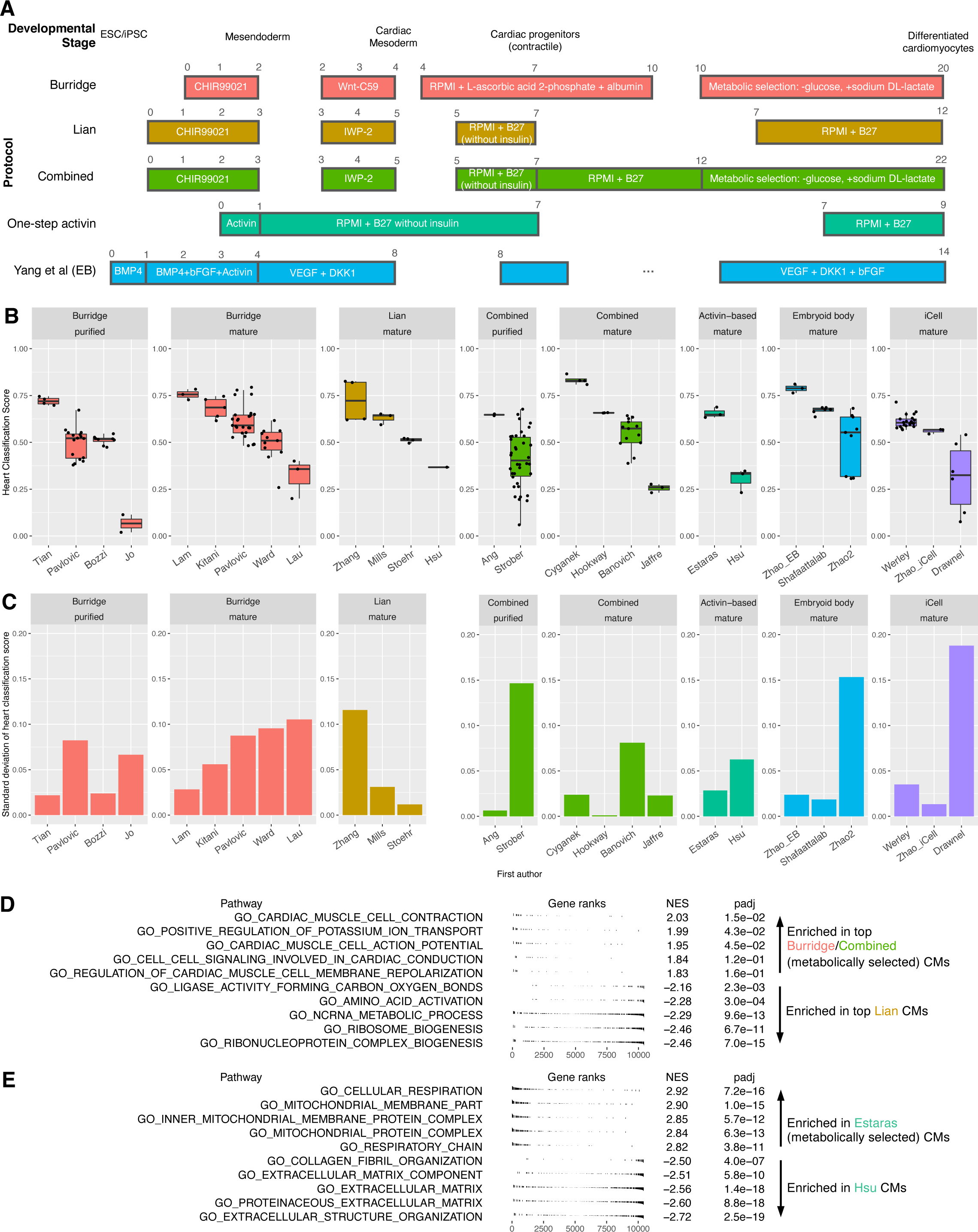
Cross-study meta-analysis of cardiomyocyte engineering protocols. See also Figures S2-5, S7. (A) Schematic of representative protocols in directed differentiation of cardiomyocytes. Protocols are aligned by developmental stage. ESC: embryonic stem cell. iPSC: induced pluripotent stem cell. EB: embryoid body. (B) Heart classification scores by study/first author and protocol for healthy/unperturbed purified and mature CM samples (from the top performing protocol variant, if relevant) per study. Within each facet, studies are ordered by decreasing mean heart classification score. (C) Standard deviation in heart classification scores by study/first author and protocol for the same purified and mature CM samples as in (B). (D) Summary plot for Gene Set Enrichment Analysis (GSEA) performed on differentially expressed genes comparing profiles of top classifying CMs from the Burridge and Combined protocols (Tian, Lam, Cyganek, Hookway, which include a metabolic selection step) versus top-classifying CMs from Lian protocol (Mills, Stoehr, which do not include a metabolic selection step). Top 2 classifying samples by Zhang were excluded from this comparison, as they were purified using a study-specific method of sorting for MYL2 positivity. Note that for this analysis and all others below that entailed examining expression of genes across studies, we re-processed all corresponding RNA-seq data using the in-house pipeline alignment and expression estimation (see Methods). (E) Summary plot for GSEA performed in differentially expressed genes between two Activin-based protocol studies: Estarás et al, who did perform metabolic selection vs Hsu, who did not. NES: normalized enrichment score. padj: Adjusted p-value based on Benjamini-Hochberg correction.

To assess the comparative transcriptional similarity of these protocols to in vivo CMs, we acquired publicly available bulk expression data generated following each method as follows: eight studies following the Burridge protocol (Bozzi et al., 2019; Jo et al., 2020; Kitani Tomoya et al., 2020; Lam Chi Keung et al., 2019; Lau et al., 2019; Pavlovic et al., 2018; Tian Lei et al., 2020; Ward and Gilad, 2019), four studies exclusively following the Lian protocol (Hsu et al., 2018; Mills et al., 2018; Stoehr et al., 2019; Zhang et al., 2019b), six studies using combined adaptations of the Lian protocol with an additional metabolic selection step (Ang et al., 2016; Banovich et al., 2018; Cyganek et al., 2018; Hookway et al., 2019; Jaffré Fabrice et al., 2019; Strober et al., 2019), two studies following a one-step activin-based differentiation protocol (Estarás et al., 2017; Hsu et al., 2018), three studies following EB protocols (Shafaattalab et al., 2019; Zhao et al., 2019), and two studies using purchased iCell CMs (Drawnel et al., 2017; Werley et al., 2017), for a total of 23 studies and 809 samples across a range of derivation stages. We queried each of these studies with PACNet classifier trained with our healthy human samples (Fig. S2A, Table S4). Comparing among developmental stages, PACNet analysis showed that most CM protocols demonstrated a gradual, consistent increase in heart classification score and decrease in ESC classification score over their differentiation time courses (Fig. S2B, S3A-B). We also considered whether our classifier was corroborated by known markers of CM fate. Across samples, we computed the correlation between heart classification scores and the expression of key CM marker genes, including TNNT2 (R^2^=0.49), TBX5 (R^2^=0.35), MYL2 (R^2^=0.73) (Fig. S4A). Heart classification correlated substantially with marker expression, but was imperfect, suggesting that the aggregate metric of classification better reflects cell identity as opposed to the expression of single marker genes.

To examine more closely the changes in cell fate over the course of differentiation, for studies that provided by-day time point labels, we correlated time point with both heart and ESC classification scores (Fig. S3B). To facilitate a fair comparison for this and subsequent analysis, we excluded any CMs with disease phenotypes, drug exposure, or genetic perturbations that might obscure accurate assessment of protocol fidelity. In cases where a study had multiple protocol variations, we selected the variant that yielded samples with the highest mean classification score for the target cell type or tissue. On average, heart classification increased with longer culture time but plateaued at approximately day 30. The single highest engineered heart classification score among all studies was achieved by day 90 ventricular CMs generated by Cyganek (Cyganek et al., 2018) (Fig. S2C), who combined aspects of Burridge and Lian during early differentiation and performed metabolic selection as described in Burridge, from day 15 to day 20. An exceptionally high score for the Lian protocol was achieved in the day 80 ESC-derived ventricular CMs generated by Zhang (Zhang et al., 2019b), who further added a step of FACS-sorting for MYL2 (myosin light chain 2v) prior to sequencing (Fig. S2C). Although the highest classifying samples were cultured for 80 and 90 days, we observed day 20 samples that achieved classification scores of the same magnitude, and day-240 samples with substantially lower classification (Fig. S3B). ESC classification decreased to negligible levels at approximately day-18 across all protocols (Fig. S3B). Thus, although increased time in culture was generally associated with a higher PACNet heart classification score, shorter protocols were still able to achieve near maximal classification scores.

We noticed that even within the same protocol, study, and time point, classification scores of replicates could vary greatly. Intra-lab and inter-lab reproducibility and consistency are important characteristics to consider in cell engineering; thus, we asked to what extent CMs engineered using the same protocol vary within and across studies. To do so, we computed the PACNet heart classification score for individual studies within and across protocols for “purified” CMs (metabolically selected CMs not yet designated as “mature”) from Burridge and Combined and all mature CMs (Fig. 2B). In one notable case, a large study that generated almost 300 samples across 19 independent rounds of differentiation (Strober et al., 2019) produced purified CMs with heart classification scores ranging from .016 to 0.611 (Fig. 2B, Fig. S2A-B). Other studies with higher variability in classification score included Ward and Pavlovich (Burridge protocol), Banovich (Combined protocol), and Zhao2 (EB protocol) (Fig. 2C). The source of this intra-study variation is unclear; however, one likely contributor is PSC line-specific differentiation bias. For example, it is known that independently derived ESC lines can generate unique differentiation biases (Abeyta et al., 2004; Bibikova et al., 2006; Osafune et al., 2008) and that iPSCs can retain epigenetic memory of their initial cell type (Kim et al., 2011). On the other end, studies with the least variability in score included Tian and Lam (Burridge protocol), Stoehr (Lian), Ang, and Cyganek (Combined protocol), Estarás, (Activin-based protocol), Shafaattalab (EB protocol), and Zhao (commercially available iCell CMs) (Fig. 2C). The studies which achieved the highest mean score within their respective protocols often achieved the most consistent scores as well, in the case of Tian, Lam, Ang, Estarás, and Shafaattalab (Fig. 2B-C). These all generated CMs from an intra-study-consistent starting cell type: Tian and Hookway from fibroblast-derived iPSCs, and Lam from PBMC-derived iPSCs. Estarás and Shafaattalab both derived CMs from ESCs. Whereas the highest mean score often achieved the most consistent scores for intra-study comparisons, *inter*-study classification scores varied more greatly for all protocols, with a standard deviation (StdDev) in heart classification score of 0.153 (Burridge) and 0.152 (Combined) for purified CMs, and ranging from 0.115 (Lian) to 0.198 (Activin-based) for mature CMs (Fig. S2D). The ability of a protocol to produce the most highly classifying CMs did not relate to protocol variability, with comparable top-scoring samples observed in the Lian, Burridge, Embryoid body, and Combined protocols irrespective of protocol StdDev.

To investigate how differences in protocols contributed to differences in heart classification, we next examined differential expression among purified and mature CM samples derived via the best performing study per protocol. We were particularly interested in the effects of metabolic selection via glucose deprivation and/or sodium DL-lactate supplementation on heart classification score, as this step was performed in 15 out of 25 studies that produced purified or mature CMs. Thus, we selected the top-classifying Burridge and Combined studies (Tian, Lam, Cyganek) to compare against the top-classifying Lian studies (Zhang, Mills). We limited the Zhang samples to the two that had not undergone additional study-specific purification via sorting for myosin light chain 2 (MYL2) positivity. Gene set enrichment analysis (GSEA) of the differentially expressed genes revealed an enrichment in gene sets related to cardiac morphogenesis and action potential in Burridge and Combined samples (Fig. 2D), corroborating an increased CM fate in response to metabolic selection. We also examined another relevant comparison within the Activin-based protocol, in which Estarás performed metabolic selection but Hsu did not. GSEA revealed an enrichment of gene sets related to cellular respiration and inner mitochondrial membrane function (oxidative phosphorylation) in the metabolically selected Estaras samples (Fig. 2E). This is consistent with the known transition from more glucose- and glycolysis-dependent metabolism during embryonic heart development to more mitochondrial- and fatty acid oxidation-dependent metabolism after birth (Chung et al., 2007; Lopaschuk and Jaswal, 2010; Piquereau and Ventura-Clapier, 2018; Tohyama et al., 2013). The higher classification scores of metabolically selected mature CMs may reflect this transition away from glycolysis and towards oxidative phosphorylation. Interestingly, we noticed that the two Zhang samples that underwent sorting for MYL2 positivity achieved heart classification scores comparable to the top-classifying Burridge and Combined studies. Thus, we asked whether this sorting increased the heart classification scores reflects a common underlying metabolic phenotype. GSEA comparing the Zhang MYL2-sorted samples to top-classifying non-sorted Lian samples demonstrated an enrichment in gene sets related to cardiac muscle function but surprisingly also related to glucose catabolism (Fig. S5A). In fact, in comparing against Burridge and Combined samples, we observed the same enrichment in glucose catabolism in the MYL2-sorted Zhang samples (Fig. S5B). This suggests that Burridge and Combined samples achieve a strong CM identity through a different mechanism compared to MYL2-sorted Zhang samples. This suggests the potential for a compounded increase in CM identity if metabolic selection is combined with sorting for MYL2.

It has been observed that both directed differentiation and direct conversion can activate the transcriptional programs of unintended cell types (Cahan et al., 2014; Kong et al., 2022; Morris et al., 2014). Therefore, we assessed whether there were any off-target lineages detectable in these engineered CMs. Although off-target effects were not prominent, we found three minor off-target signatures among CM studies. Two studies showed a noticeable aberrant fibroblast signature, though the mature CM samples predominantly still classified as heart (Fig. 3A). We corroborated the fibroblast signature with key markers of fibroblast identity, including COL1A1 and THY1 (Fig. 3B). The latter of the two studies, Hsu et al (Hsu et al., 2018), also showed a noticeable intestinal signature (Fig. 3A), which we corroborated with expression of key intestinal genes, including SLC10A2 and MUC2 (Fig. S6A). Interestingly, the metabolically selected counterparts (Estarás et al., 2017) to the Hsu CMs lacked both these off-target signatures (Fig. S6B), suggesting that metabolic selection may act to remove off-target cells as well as other cardiac lineage cells (Andersen et al., 2018; Zhang et al., 2019a). A subset of samples from two studies (from the same publication (Zhao et al., 2019)) demonstrated an off-target liver signature (Fig. 3C), which we corroborated by examining expression of key liver markers including ALB, SERPINA1, and CEBPA (Fig. 3D). These two studies followed EB protocols for CM differentiation, followed by seeding onto tissue biowire constructs and application of electrical stimulation (Zhao et al., 2019). It is worth noting that early activation of Activin/Wnt signaling also specifies the definitive endoderm adjacent to precardiac mesoderm (Kubo et al., 2004; Toivonen et al., 2013) and endodermal derivatives are often present in cardiac organoids (Drakhlis et al., 2021; Rossi et al., 2021). Thus, these protocols may be permissive to hepatic differentiation as well.

**Figure 3.**
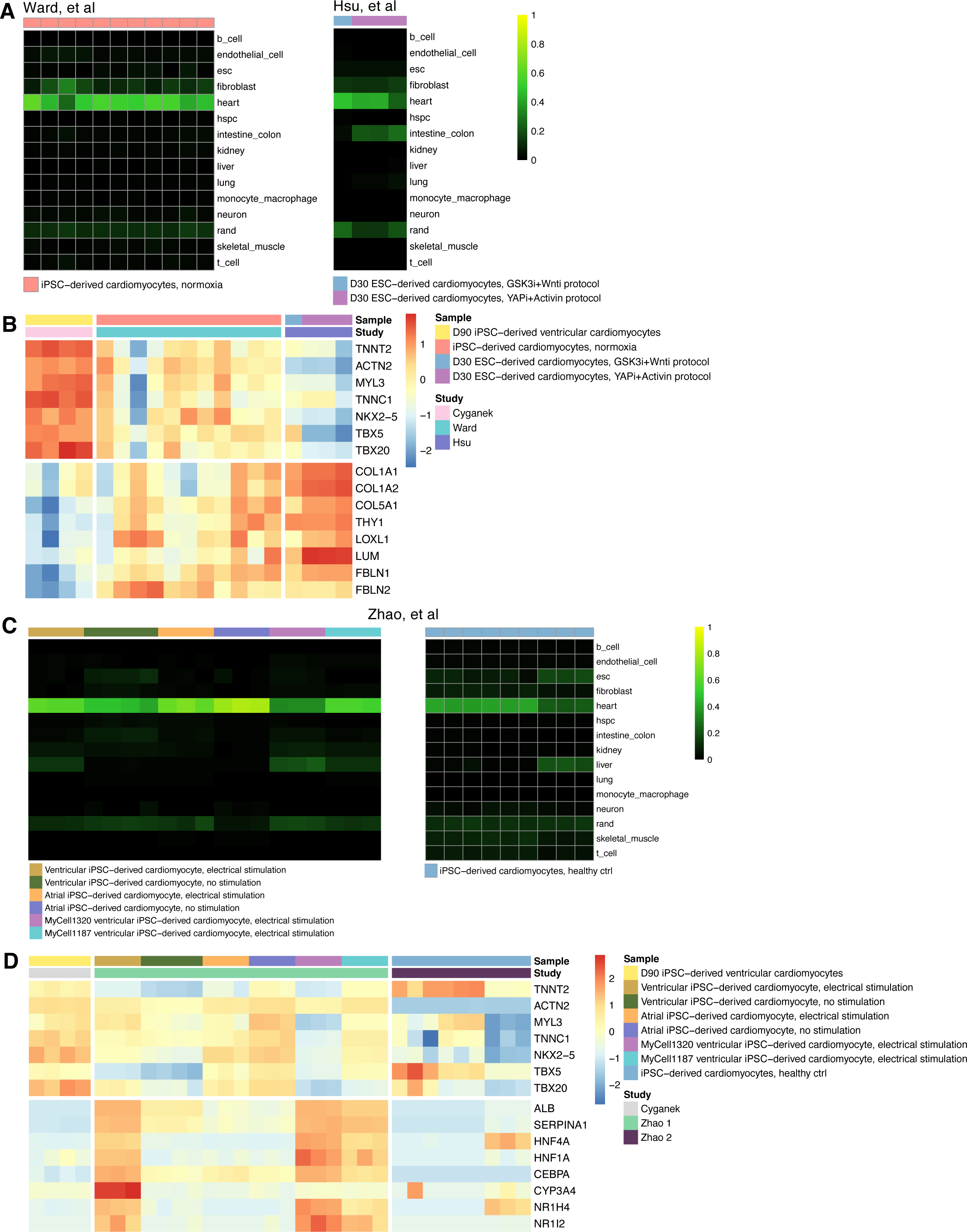
Off-target signatures in cardiomyocyte engineering. See also Figure S6. (A) Heatmap of classification scores for Ward et al (left) and Hsu et al (right) studies which produced cardiomyocytes with an off-target fibroblast signature. (B) Heatmap of gene expression for canonical cardiomyocyte and fibroblast marker genes for the study with the most highly classifying cardiomyocytes (Cyganek et al) compared to Ward et al and Hsu et al studies. (C) Heatmap of classification scores for two separate GEO accession studies from the same authors (Zhao et al) which produced cardiomyocytes with an off-target liver signature. Heatmap color scale: z-score of log-transformed gene counts. (D) Heatmap of gene expression for canonical cardiomyocyte and liver marker genes for the study with the most highly classifying cardiomyocytes (Cyganek et al) compared to Zhao et al studies. For B and D: Heatmap color scale: per gene z-score of log-transformed TPM.

Finally, we sought to identify transcription factors (TFs) whose modulation might improve engineering protocol performance. To do so, we used an adapted version of the CellNet Network Influence Score (Cahan et al., 2014) (NIS, see Methods), which evaluates the need for up- or down-regulation of cell type-specific TFs based on the expression of those TFs and their target genes. Across engineered mature CMs, NKX2-5 (NK2 Homeobox 5) was most strikingly assigned the highest mean score, indicating a predicted need for upregulation. This is consistent with the role of NKX2-5 as a master regulator of CM fate, controlling a subnetwork of CM transcription factors including TBX5, TBX20, HAND1, and HAND2 (Akazawa and Komuro, 2005), which also had moderately strong positive scores in return (Fig. S7A). The NIS successfully prioritized the upregulation of NKX2-5 in CM samples with more prominent off-target signatures (Fig. S7A), including Ward (with off-target fibroblast, Fig. 3A). It also recognized the already prominent NKX2-5 activity in highly classifying CM samples, including Cyganek and Zhang (Fig. 2B, Fig. S7B). Taken together, these results demonstrate the utility of PACNet in assessing cross-study performance and identifying actionable points for protocol improvement.

### Hepatocyte engineering studies have diverse performance outcomes

We next identified common hepatocyte derivation protocols in the field. With less consensus in specific derivation protocols than seen with cardiomyocytes, we divided hepatic protocols into four general categories: directed differentiation (henceforth denoted as DD), ESC/iPSC-derived hepatic organoid differentiation (PSC-O), transdifferentiation (TD, i.e. direct conversion), and primary hepatocyte-derived organoid (P-O) culture. We analyzed 24 studies, divided as follows: twelve DD studies (Ang et al., 2018; Boon et al., 2020; Carpentier et al., 2020; Deaton et al., 2018; Du et al., 2014; Ehrlich et al.; Gao et al., 2017; Qin et al., 2016; Sagi et al., 2019; Tilson et al., 2021; Touboul et al., 2016; Viiri et al., 2019), six PSC-O studies (Akbari et al., 2019; Asai et al., 2017; Guan et al., 2020; Ouchi et al., 2019; Velazquez et al., 2021; Wang et al., 2019), four TD studies (Du et al., 2014; Gao et al., 2017; Wang et al., 2016; Xie et al., 2019), and eight P-O studies (Artegiani et al., 2019; Broutier et al., 2017; Giobbe et al., 2019; Hu et al., 2018; McCarron et al., 2021; Schneeberger et al., 2020; Wang et al., 2019; Ye et al., 2020). We summarize each of these derivation strategies below (and illustrate a representative protocol for each in Fig. 4A):

(1) DD protocols followed an overall consistent pattern, beginning with unanimous induction of definitive endoderm with Activin A, with some adding Wnt3a or BMP4. Inducing the transitions from definitive endoderm to foregut or hepatic endoderm followed two major strategies: application of BMP4 and FGF2 (Boon et al., 2020; Deaton et al., 2018; Qin et al., 2016) or of 2-mercaptoethanol and DMSO (Carpentier et al., 2020; Ehrlich et al.; Sagi et al., 2019). Other strategies included BMP4 combined with TGFβ inhibition and Wnt inhibition (Touboul et al., 2016). Differentiation to hepatoblasts and hepatocytes or hepatocyte-like cells was, with large consensus, induced with some combination of Oncostatin M, Dexamethasone, and hepatocyte growth factor (HGF).
(2) PSC-O protocols followed global strategies similar to DD, but with the addition of various common organoid-promoting factors including R-spondin1, EGF, Noggin, FGF10, and gastrin (Huch et al., 2013; Pleguezuelos-Manzano et al., 2020) at various stages. Two studies incorporated TGFβ inhibitor SB431542 and adenylyl cyclase activator forskolin (Akbari et al., 2019; Wang et al., 2019) during specification to mature hepatic organoids, with one distinct study also adding BMP7 and Notch signaling inhibitor DAPT during this final stage (Akbari et al., 2019). A unique organoid protocol achieved differentiation through transduction of transcriptional regulators (TFs) GATA6, PROX1, and ATF5, as well as CRISPR-mediated activation of endogenous enzyme CYP3A4 (Velazquez et al., 2021).
(3) Transdifferentiation of fibroblasts into hepatocytes involved transduction of specified hepatic TFs, most commonly a combination of HNF family and FOXA family TFs, followed by culture in hepatocyte induction media (Du et al., 2014; Gao et al., 2017; Xie et al., 2019). One unique transdifferentiation method reverted gastric epithelial cells back to an endodermal progenitor state using a cocktail of small molecule inhibitors, followed by a standard hepatocyte differentiation procedure (Wang et al., 2016).
(4) In P-Os, two major protocols, which we hereafter denote by first author, were used: Huch et (Huch et al., 2015) and Hu (Hu et al., 2018). Huch established a system to expand adult bile duct-derived bipotent liver progenitor organoids, which upon supplementation with additional factors, can give rise hepatocytes. Hu then elaborated on this system to establish long-term expansion conditions specifically for hepatocyte-derived organoids.

**Figure 4.**
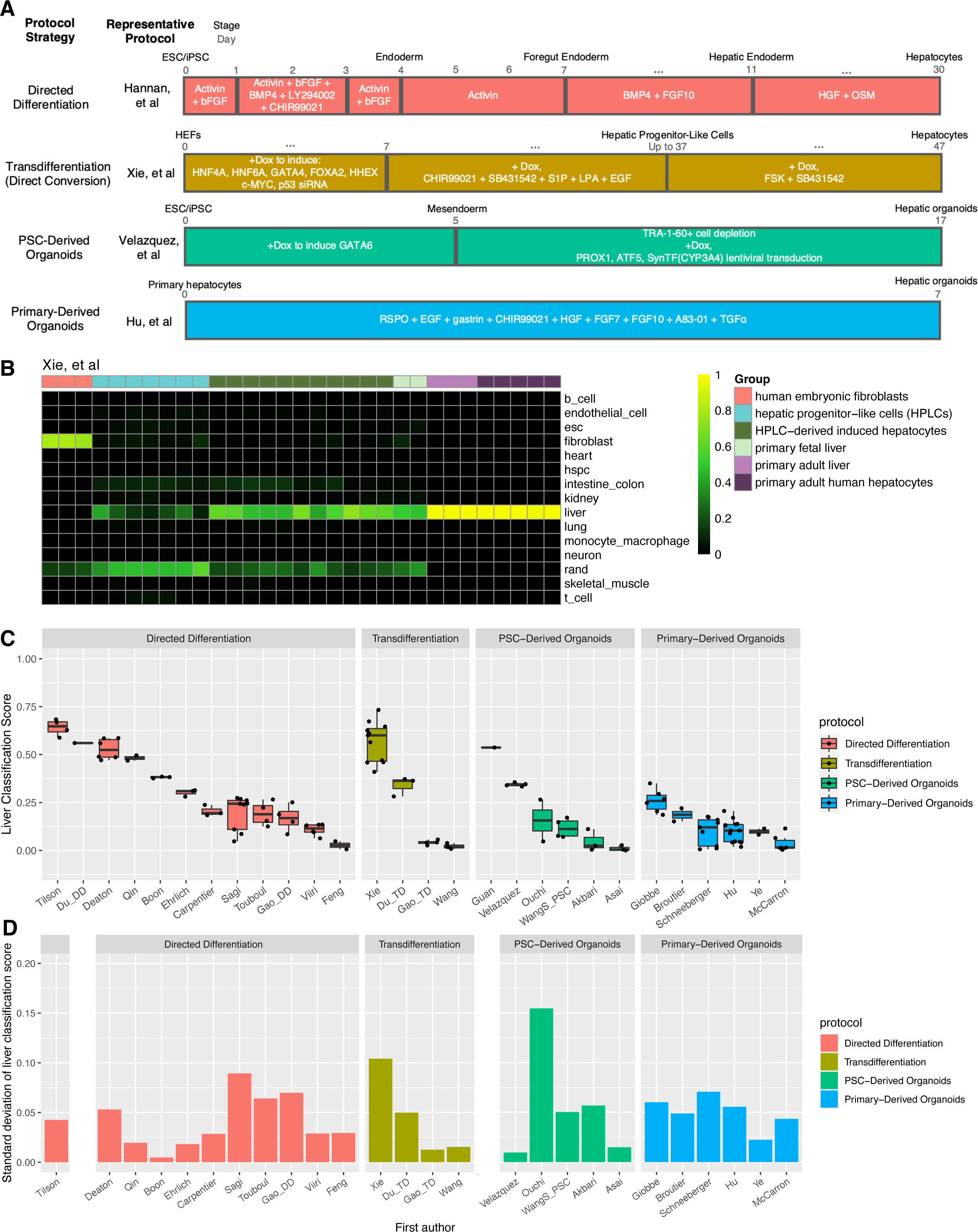
Cross-study meta-analysis of hepatocyte engineering protocols. See also Figures S8-S12. (A) Schematic of representative protocols from each hepatocyte derivation strategy. ESC: embryonic stem cell. iPSC: induced pluripotent stem cell. HEF: human embryonic fibroblast. SynTF(CYP3A4): synthetic transcription factor for endogenous CYP3A4 activation. (B) Classification heatmap for study that produced the most highly classifying hepatocyte samples (Xie et al transdifferentiation from human embryonic fibroblasts to hepatocytes). (C) Liver classification scores by study/first author and protocol for healthy/unperturbed, fully mature or differentiated hepatocyte and hepatic organoid samples (from the the top-performing protocol variant, if relevant) per study. Within each facet, studies are ordered by decreasing mean liver classification score. (D) Standard deviation in liver classification scores by study/first author and protocol for the same hepatocyte or hepatic organoid samples as in (C).

We first assessed the transcriptional similarity of these studies to in vivo hepatocytes using PACNet (Fig. S8A). Whereas engineered cardiomyocyte showed a consistent gradual increase in heart classification score over the course of differentiation, engineered hepatocytes generally experienced a late, sharp increase in liver classification score in the transition from hepatic progenitor to mature hepatocytes (Fig. S8B-C). Studies demonstrated a concurrent decrease in ESC classification score (or fibroblast classification score for relevant transdifferentiation studies), though most samples retained a detectable ESC signature (Fig. S9A-C). We also assessed whether the classifier was corroborated by known markers of hepatocyte identity. Though on average classification score increased with increasing expression of hepatocyte marker genes, correlation coefficients were low among all engineered samples: ALB (R^2^=0.33), SERPINA1 (R^2^=0.35), and HNF4A (R^2^=0.11) (Fig. S9D), again suggesting that the aggregate metric of classification better reflects cell identity as opposed to the expression of individual marker genes Next, we again asked which specific protocols produced populations with the highest PACNet classification score. The highest classifying mature hepatocytes were produced by Xie et al (Xie et al., 2019) (Fig. 4B-C), who used a two-step, 37 to 47-day TD protocol. This protocol first converts embryonic fibroblasts into hepatic progenitor cells through the transduction of HNF4A, HNF6A, GATA4, FOXA2, and HHEX, followed by differentiation into functionally competent mature hepatocytes using forskolin and ALK5 inhibitor SB431542. Almost equally well-classifying were two DD studies performed by Tilson (Tilson et al., 2021) and Deaton (Deaton et al., 2018) (Fig. 4C, S10A-B). Tilson and Deaton each performed adaptations of a protocol by Hannan (Hannan et al., 2013): a 20-day differentiation through primitive streak, definitive endoderm, and hepatic endoderm states to generate mature hepatocyte-like cells. While Tilson followed the Hannan method of including PI3K inhibitor LY294002 during definitive endoderm induction, Deaton used a commercially available definitive endoderm kit. Deaton also uniquely included myosin II inhibitor blebbistatin during the induction of hepatic endoderm from definitive endoderm. Finally, a DD protocol variation performed by Qin that included supplementation of Vitamin K2, known to contribute to hepatocyte maturation (Avior et al., 2015b) and to increase expression of mature hepatocyte gap junction protein Connexin 32 (Qin et al., 2016), also had one of the highest mean classification scores (Fig. 4C, S10C).

We next examined intra- and inter-study variability in classification score and their relationship to study performance. To facilitate a fair comparison, we again excluded any engineered samples with disease phenotypes, drug administrations, or genetic perturbations that were unrelated to protocol improvement. And if a study had multiple protocol variations, we selected the variation producing the more highly classifying samples (by mean). In CMs, we had observed that intra-study consistency increased with higher mean heart classification score. In contrast, for hepatocyte derivation overall, StdDev was not associated with mean PACNet classification and did not reflect high protocol performance. (Fig. S10D, Fig. 4C-D). Only the aforementioned Tilson DD study had a mean classification of over 0.5 while maintaining a StdDev under 0.05. The Xie study, which did have a mean classification score of over 0.5, had a higher StdDev of 0.089. Of the PSC-O and P-O studies, Velazquez and Giobbe, respectively, had the highest mean classification scores while maintaining StdDevs under 0.05 (Fig. 4C-D). At the inter-study level, while DD and TD protocols had higher StdDevs of 0.259 and 0.190 respectively, DD and TD samples significantly outperformed P-O and PSC-O samples on average (Fig. S8D). The extent to which inter-study and inter-protocol variability in PACNet score predict or reflect functional variability in engineered populations is unknown. Nonetheless, having a quantitative metric of population quality will be useful as the field places more emphasis on reducing the variability within protocols.

We next wanted to investigate to what extent DD and TD protocols might each have independent qualities that could help improve the other. To this end, we performed differential expression and gene set enrichment analysis (GSEA) on mature samples from the most highly classifying DD study (Tilson) and the most highly classifying transdifferentiation study (Xie). GSEA revealed in Xie-derived hepatocytes a slight enrichment of genes involved in liver-related functions, including fatty acid metabolism (CYP4A11, EHHADH) and bile acid metabolism (NR1I2, NR1H4, SLC27A5). On the other hand, the Tilson-derived hepatocytes demonstrated a slight enrichment of genes (including PROS1, FGG, FGA, F9) related to the blood coagulation system, in which hepatocytes are known to play a role (Kopec and Luyendyk, 2014) (Fig. S11A-B). Therefore, approaches that leverage aspects of both protocols could lead to additive increases in hepatocyte identity.

To determine other potential enhancements to these protocols, we again sought to identify liver-specific TFs whose modulation might improve classification performance, using the adapted CellNet NIS. In mature engineered hepatocytes, PROX1 (Prospero Homeobox 1) and NR1H4 (Nuclear Receptor Subfamily 1 Group H Member 4, also known as Farnesoid X Receptor) scored most highly across a variety of protocols, consistent with their roles as key hepatic regulators (Preidis et al., 2017; Seth et al., 2014) (Fig. S12A). The NIS prioritized upregulation of NR1H4 in engineered samples with modest liver PACNet scores and prominent off-target PACNet scores, including from Sagi and Feng (with high residual ESC signatures, Fig. S10E-F), from Hu (off-target neuronal signature, Fig. 6A-B), from Wang (off-target fibroblast signature, Fig. S13A), and from Asai (with off-target endothelial and fibroblast signatures, Fig. S13A). Furthermore, this analysis recognized that sufficient NR1H4 activity was present in highly classifying hepatocyte samples generated by Tilson (Fig. S10A), Qin, (Fig. S10C), and Xie (Fig. 4B). Specifically examining the expression of NR1H4 targets, Tilson, Qin, and Xie samples indeed had higher expression of key hepatocyte genes, including ALB, APOA1, and APOC3 (Fig. S12B). Thus, PACNet successfully exploited its cross-study capabilities to characterize the global performance of engineered liver studies.

We finally explored off-target signatures in engineered hepatocytes, which were more prevalent than in CM engineering studies (Fig. S8A, S2A). PACNet detected a consistent aberrant raised intestine/colon classification score in several stem cell-derived and primary liver-derived organoid studies (Schneeberger, Ouchi, McCarron, Broutier, Akbari, Ye, WangS, Fig. 5A). This was unsurprising given the shared endodermal lineage of intestine and liver and because of shared culture medium components with intestinal organoid culture protocols, including R-spondin1 and EGF (Pleguezuelos-Manzano et al., 2020). Expression of intestinal marker genes including CDX1, CDX2, and SI corroborated the off-target classification (Fig. 5B). Interestingly, the samples derived from different studies seemed to express some mutually exclusive intestinal markers, suggesting further specification into intestinal cell subtypes. Upon examination of specific subtype markers, most studies showed some enterocyte and goblet cell lineage enrichment, with McCarron samples being most consistently enriched in these markers. Ouchi samples had a slight tuft cell enrichment based on expression of markers DCLK1 and NREP, while Akbari and Ye samples had increased expression of both enteroendocrine (CHGA, NEUROD1, NKX2−2) and tuft cell markers (Fig. 5B). This intriguingly suggests potential avenues for intestinal subtype-specific cell derivation protocols.

**Figure 5.**
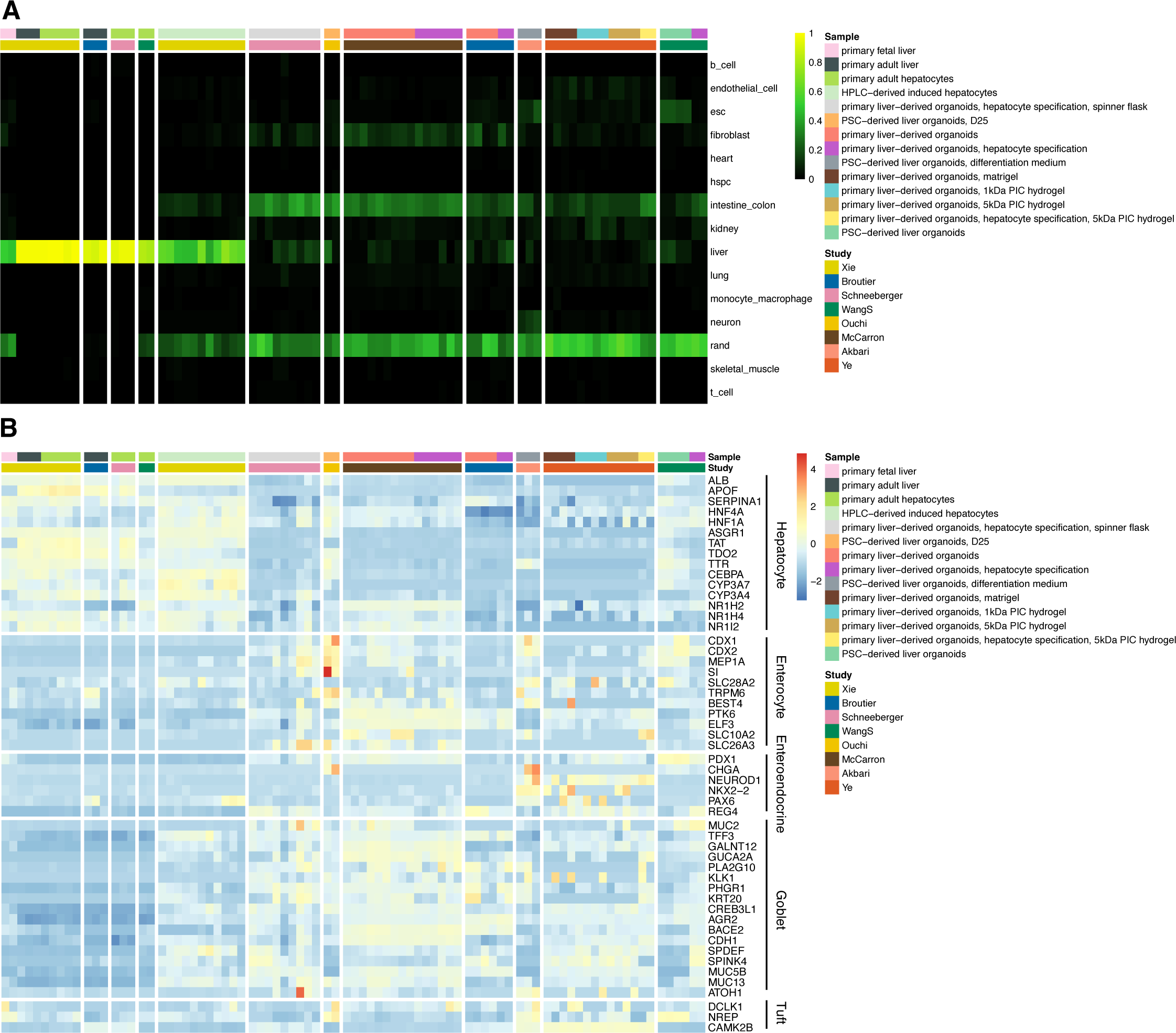
Off-target intestinal signatures in stem cell-derived hepatic organoids. See also Figure S13. (A) Classification heatmaps for highest classifying hepatocyte derivation study (Xie et al) vs Broutier et al, Schneeberger et al, Wang S et al, Ouchi et al, McCarron et al, Akbari et al, and Ye et al. (B) Heatmap of hepatocyte and intestinal (split into subcategories) marker gene expression for the same studies as in (A). Heatmap color scale: per gene z-score of log-transformed TPM.

**Figure 6.**
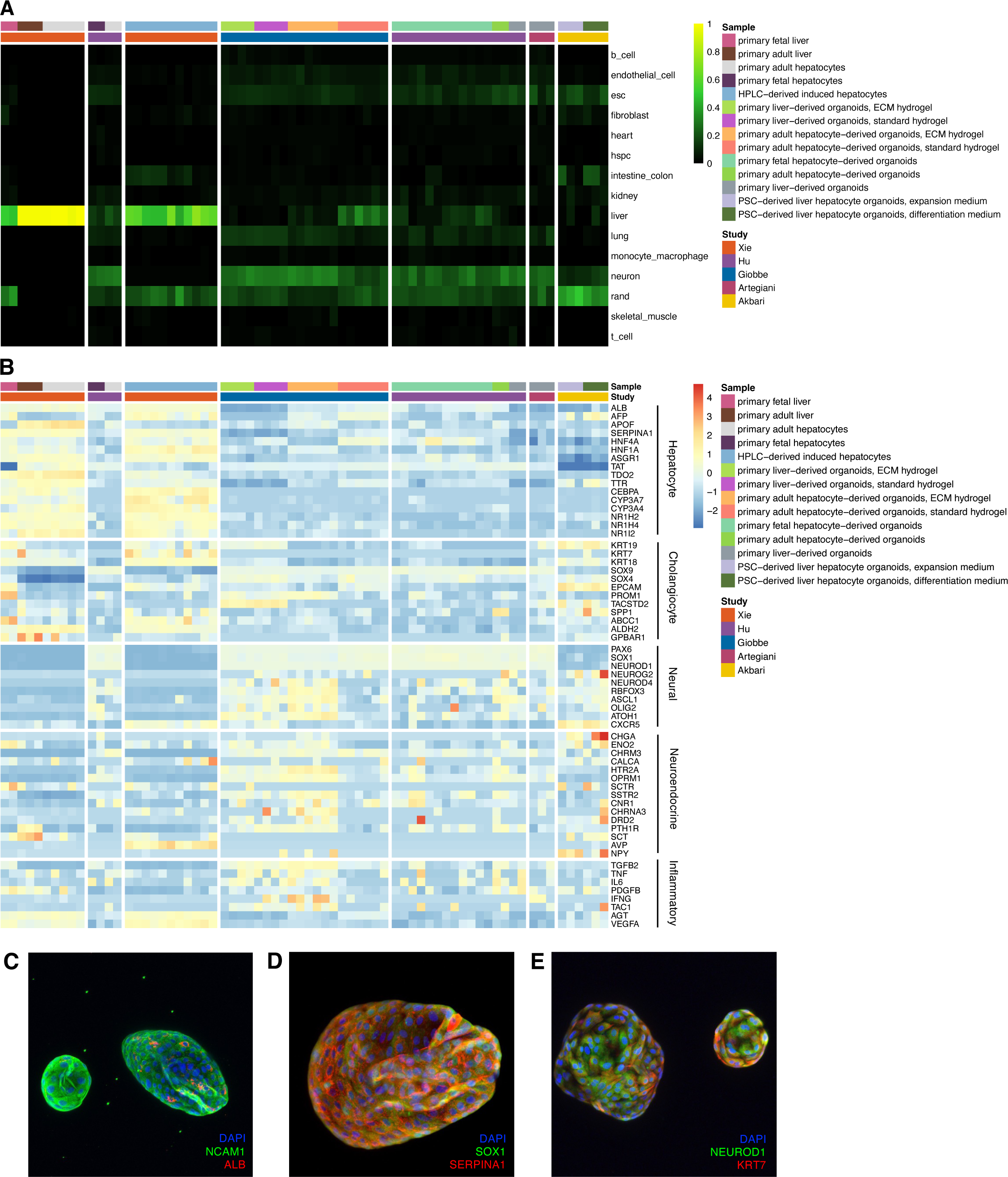
Off-target neuronal signatures in primary hepatocyte-derived organoids. See also Figure S14-15. (A) Classification heatmaps for highest classifying hepatocyte derivation study (Xie et al) vs Hu et al, Giobbe et al, Artegiani et al, and Akbari et al. (B) Heatmap of hepatocyte and neuronal marker gene expression for the same studies as in (A). Heatmap color scale: per gene z-score of log-transformed TPM. Immunofluorescence staining for (C) neuroendocrine marker NCAM1 (green) and hepatocyte marker Albumin (ALB, red); (D) neuronal marker SOX1 (green) and hepatocyte marker SERPINA1 (red); and (E) neuronal marker NEUROD1 (green) and cholangiocyte marker KRT7 (red). Blue: DAPI.

We also observed a fibroblast signature in gastric epithelial cell-derived hepatocytes derived by Wang (Wang et al., 2016) and both an endothelial cell and fibroblast signature in iPSC-derived liver bud cells derived by Asai (Asai et al., 2017) (Fig. S13A). In the former study, though the derived cells expressed hepatocyte markers APOF and NR1H2 at moderate levels, the expression of fibroblast markers including PDGFRA and FBLN2 dominated (Fig. S13B). The off-target signature may reflect a carryover of stromal cells from co-culture with gastric subepithelial myofibroblast feeder cells at the endodermal progenitor stage (Wang et al., 2016). In the latter, though there was some expression of liver markers in the derived liver bud organoids (Fig. S13B), the moderate endothelial classification signature and the strong expression of fibroblast marker genes most likely reflect co-culture of hepatic-specified endoderm cells with human umbilical vein endothelial cells and mesenchymal stem cells during organoid formation (Asai et al., 2017). The successful identification of known co-culture cell types further corroborates the ability of PACNet to identify non-physiological levels of off-target cell type signatures.

### Primary liver-derived organoids exhibit consistent off-target neural signatures

Surprisingly, we observed that several studies of P-Os and one PSC-O study - Hu, Giobbe, Artegiani, and Akbari - showed aberrant neuronal classification (Fig. 6A). This was unexpected, as the nervous system and liver do not share a common lineage background (Gordillo et al., 2015), and the healthy human liver does not exhibit prominent neuronal signatures (Fig. 1A, 5A, S10F). Thus, we more closely examined what genes might contribute to this off-target signal. We confirmed expression both of canonical neural marker genes including PAX6, SOX1, NEUROD1, NEUROD4, and ASCL1 (Fig. 6B) and of key neuronal genes from the PACNet classifier (Fig. S14A, Table S11) in samples which had neuronal classification. To further investigate this signature and avoid potential batch effects that might arise when integrating data from multiple studies, we more closely examined Giobbe et al, in which hepatocyte organoids cultured in “standard hydrogel” had a stronger liver score and weaker neural score, while hepatocyte organoids cultured in “ECM hydrogel” had a stronger neural score and a negligble liver score (Fig. S14B). GSEA on genes differentially expressed between hepatocyte organoids cultured in ECM vs standard hydrogel identified neocortex gene sets in the former, which include ASCL1, NEUROD4, and SLC17A6, and an enrichment of liver-specific signatures in the latter, which include ALB, SERPINA1, and HNF4A (Fig. S14C-D). The independent detection of brain-related genes sets by GSEA in samples which had a neuronal PACNet classification further corroborates our observation of aberrant neural transcriptional programs in some hepatocyte organoids. We noted that the primary adult hepatocyte and primary fetal hepatocyte positive control samples in the Hu study also showed aberrant neuronal classification and expression of neural marker genes (Fig. 6A-B), potentially suggesting a technical issue with the classifier. However, PACNet successfully classified primary fetal liver, adult liver, fetal hepatocyte, and adult hepatocyte samples from many other studies (Asai, Boon, Du, Gao, Qin, Schneeberger, Touboul, Viiri, WangS, and Xie) as liver and not neural (Fig. S8B, S15A). Therefore, the aberrant neural signature seen in these primary samples from the Hu study are unlikely to be due to classifier performance, but rather potentially arose from mis-annotation of the samples (i.e. some organoid samples might have been inadvertently labeled as ‘primary’). Finally, we note that not all primary liver-derived organoid samples had this same neuronal signature: primary-derived organoid samples generated by Schneeberger, Ouchi, McCarron, Broutier, Ye, and WangS also lacked an aberrant neuronal signature (Fig. 5A). This prompted us to look for differences in derivation strategies that might explain the origin of the neural signature.

Primary liver-derived hepatocyte organoids fell into two broad protocols: 1) derivation of bipotent progenitor organoids from EPCAM+ cholangiocytes (WangS, Giobbe subset), followed by a hepatocyte specification step (Huch et al., 2015) (Schneeberger, Ye, McCarron) and 2) direct derivation of organoids from hepatocytes (Hu et al., 2018) (Hu, Giobbe subset). The former category also included a variation where bipotent progenitor organoids were initially supplemented with Wnt-conditioned media and Noggin (Artegiani), followed by the same hepatocyte specification step (Broutier). We have summarized these protocol differences in Figure S15B. The presence of a neural signature in only Artegiani, Hu, and the Giobbe direct hepatocyte subset potentially implicates two mechanisms of origin: 1) culture conditions specific to the Wnt-conditioned medium- and noggin-supplemented bipotent progenitor organoid stage or 2) culture conditions specific to the Hu protocol. Further studies will be necessary to identify the exact determinants of this signature.

We next investigated whether the observed neural signature was connected to related lineages with known liver roles. Neuroendocrine factors are known to be expressed during ductular reaction in cholangiopathies (Alvaro et al., 2007; Banales et al., 2019; Ehrlich et al., 2018; Munshi et al., 2011). We hypothesized that neuroendocrine gene expression might contribute to the detected neuronal signature and indeed observed that the aberrant samples also consistently expressed high levels of several neuroendocrine genes and neuropeptides, including CHGA, CHRM3, and PTH1R, relative to the most highly classifying hepatocyte samples generated by Xie (Fig. 6B). Liver ductal organoids derived by Giobbe and PSC-derived hepatocyte organoids derived by Akbari also expressed high levels of cholangiocyte or biliary marker genes, including KRT19, EPCAM, PROM1, and TACSTD2. Furthermore, cholangiocyte marker SOX9 was also moderately expressed across most aberrant neuronal samples (though SOX9 is also a broad marker of neural stem cells (Scott et al., 2010), ependymal cells (Pompolo and Harley, 2001), astrocytes (Sun et al., 2017), and intestinal cells (Blache et al., 2004)). Interestingly, although they had the lowest expression in neural and neuroendocrine marker genes, Xie engineered hepatocytes had the highest expression of cholangiocyte markers, including KRT18 and KRT7. We also examined inflammatory markers indicative of ductular reaction, including TGFB2 and IL6 (Ehrlich et al., 2018; Munshi et al., 2011; Park, 2012), and found that their expression was indeed generally associated with neural and neuroendocrine gene expression, particularly in Giobbe ductal organoids and standard hydrogel hepatocyte organoids. The fact that many neurally classifying organoid samples did not express cholangiocyte markers (and that many Xie non-neurally classifying samples expressed cholangiocyte markers) suggests that ductular reaction may be insufficient to fully explain the observed neuronal and neuroendocrine signatures. Nonetheless, ductular reaction may contribute to these signatures given that within the organoid samples, many with the highest neural and neuroendocrine expression (Giobbe ductal organoids and Akbari PSC-derived hepatocyte organoids) also had the highest cholangiocyte marker expression.

To experimentally test whether the aberrant neural and neuroendocrine signature in cultured primary liver-derived organoids can be recapitulated, we generated organoids from primary human liver using established protocols (Broutier et al., 2016; Huch et al., 2015) (Fig. S16A). We performed immunofluorescence staining for hepatic markers ALB and SERPINA1, cholangiocyte marker KRT7, neuroendocrine marker NCAM1 and neural markers SOX1, and NEUROD1. We observed very little ALB expression (Fig. 6C, Fig. S16B), consistent with a lack of mature hepatocyte identity, while the early hepatic marker SERPINA1 was more abundant (Fig. 6D, Fig. S16C). KRT7 was detectable (Fig. 6E, Fig. S16D), consistent with a ductal-like biliary phenotype (Huch et al., 2015). Positive staining of NCAM1, SOX1 and NEUROD1 corroborated the accumulated RNA-seq data of liver-derived organoids (Fig. 6C-E, Fig. S16C-E) and confirmed the presence of a neural/neuroendocrine signature. Although NEUROD1 expression was primarily localized to the cytoplasm, import of NEUROD1 to the nucleus has known regulatory dependencies, including dimerization with partner proteins (Mehmood et al., 2009) or glucose-dependent phosphorylation (Andrali et al., 2007; Wong et al.). Taken together, these data support a prominent off-target neural and neuroendocrine signature in some primary liver-derived organoids.

## Discussion

Our comprehensive analysis of major protocols in cardiomyocyte and hepatocyte engineering resulted in several observations that are relevant to the CFE field. In cardiomyocyte engineering, we found that two methods of purification at the later stages of differentiation proved especially effective in achieving the highest heart classification scores: 1) metabolic selection via glucose deprivation and sodium DL-lactate supplementation and 2) cell sorting for Myosin light chain 2 positivity. These independent purification methods were non-redundant, as differentiated MLC2v+ CMs retained a glucose catabolism signature more similar to the metabolic profile of CMs that did not undergo metabolic selection, suggesting that a combination of the two techniques could achieve even higher classification scores.

In examining variability within and across CM derivation studies, we observed that studies with higher mean classification scores generally achieved a higher consistency (via lower intra-study standard deviation). At the inter-study level, the ability of a protocol to produce the most highly classifying CMs did not relate to protocol variability, with comparable top-scoring samples observed across protocols irrespective to StdDev. In contrast, across hepatocyte and hepatic organoid engineering studies, at both the intra-lab and inter-lab levels, low variability was not associated with low PACNet scores. Our broad assessment of transcriptional fidelity of cell fate engineering protocols was not designed to determine the contributors to intra- and inter-study variability. However, on the CM side, protocols with high intra-study variability tended to have PSC originating from a range of sources, including dermal fibroblasts, mesenchymal stem cells, and peripheral blood mononuclear cells. Further investigation of cell of origin and other potential contributors to protocol variability will facilitate greater consistency in CFE protocols.

Our PACNet analysis also identified several recurring off-target or unexpected signatures in both CM and hepatocyte engineering studies. Most notably, we observed neural and neuroendocrine signatures in primary liver- and primary-hepatocyte derived organoids. We confirmed the expression of key liver, neural, and neuroendocrine marker genes both via quantitative analysis of expression profiles from the original studies, and by immunofluorescence staining of liver organoids that we generated following the same protocol. We postulate that cholangiocyte-related ductular reaction may contribute to the neural and neuroendocrine transcriptional programs that are detected in these organoids. Enrichment of neuron development gene sets has also been observed in primary murine liver organoids (Aloia et al., 2019), consistent with a cholangiocyte-neural signature relationship. Off-target cell fate signatures could also potentially be explained by lineage biases induced by unanticipated impact of added growth factors and small molecule inhibitors. Ectopic organoid culture-specific signatures have precedents in other *in vitro* contexts as well. For example, endoplasmic reticulum stress and glycolytic stress signatures are known to appear in brain organoids which are not reflected in primary fetal tissue signatures (Bhaduri et al., 2020; Vértesy et al., 2022). Human kidney organoid culture protocols are also known to exhibit unexpected neural signatures (Howden et al., 2019; Liu et al., 2020; Wu et al., 2018b), potentially supporting an organoid culture-specific artifactual neural signature. Further work will be necessary to determine specific mechanisms that induce off-target gene expression signatures in organoids and to characterize shared aberrant patterns of intestinal, neural, and other signatures across organoid derivation protocols, which could have broad implications for organoid derivation, maintenance, and their applications.

We note several caveats in this study and areas for improving the PACNet platform. First, as PACNet takes as input bulk RNA-Seq data for both training and query, it is unable to distinguish hybrid signatures or states from population heterogeneity in query engineered samples. As more single-cell RNA-Seq studies are performed across diverse cell engineering protocols for a variety of cell types and across corresponding reference data types (Almanzar et al., 2020; Schaum et al., 2018), we can extend PACNet to provide comparisons of single-cell transcriptional studies in cell engineering. Nonetheless, our bulk analysis still accurately evaluates cross-sample and cross-study performance and detects off-target signatures, whether they arise from hybrid states or population heterogeneity. Second, PACNet analysis is limited to transcriptional data and does not consider epigenetic, proteomic, or functional characteristics. Transcriptional profiles have modest correlation with proteomic (Abreu et al., 2009; Wang et al., 2014) and epigenetic (Jjingo et al., 2012; Starks et al., 2019; de la Torre-Ubieta et al., 2018) profiles in humans. Future efforts will be necessary to see if integration of proteomic and epigenetic data types, such as ATAC-Seq and whole-genome bisulfite sequencing, can further improve PACNet performance. Finally, PACNet was trained using samples from adults and thus this platform is likely to be less sensitive to engineered populations that are equivalent to embryonic or fetal stages of development. As more transcriptomic data accrue of developmental stages, especially using single cell and single nucleus RNA-seq, we envision that classifiers that predict developmental stage of specific tissues and lineages will be readily trained and applied.

We provide PACNet as both a web platform (http://cahanlab.org/resources/agnosticCellNet_web/) through which users can browse engineered reference panels or perform analysis, and as code that can be downloaded and executed locally, as well as modified (https://github.com/pcahan1/CellNet). PACNet provides highly precise and sensitive classification to detect on- and off-target transcriptional signatures and makes predictions for improved expression of cell/tissue type-specific transcriptional regulators.

Importantly, PACNet also facilitates comparison to current state-of-the-field engineering protocols for seven major cell and tissue types: liver, heart, neuron, skeletal muscle, lung, intestine/colon, and hematopoietic stem and progenitor cells. Thus, we think that PACNet will be a useful, unbiased tool to quantify the performance of cell fate engineering protocols and as valuable resource to the cell engineering community.

### Experimental Procedures

#### RNA-Seq preprocessing and adapting author-provided expression data

Publicly available RNA-sequencing and expression microarray datasets for query and training were curated from the NCBI Gene Expression Omnibus (GEO) (Table S1). For studies not providing preprocessed gene-by-sample expression matrices, we used the following preprocessing pipeline as previously described (Radley et al., 2017). First, adapters and low-quality bases were trimmed from FASTQ files using cutadapt. Trimmed FASTQs were mapped and quantified using salmon in mapping-based mode. For quality control, 1% of reads were randomly sampled and aligned to *E. coli*, *S. cerevisiae*, *D. rerio*, *M. musculus*, and *H. sapiens* genomes. Reads aligning to human genome features, including rRNA, mtRNA, and protein coding, were computed using HTSeq. For analyses that entailed examining expression of genes across studies (Fig 2D, Fig 2E, Fig 3B, Fig 3D, Fig 5B, Fig 6B,SFig 5A-B, SFig 6A, SFig 7B, SFig 11A-B, SFig 12B, SFig 13B, SFig 14A,C-D), we preprocessed all samples using the above pipeline.

### Cell/tissue-specific classifier training and validation

Primary human bulk RNA-sequencing training data was acquired from NCBI GEO for the following cell and tissue types: B cells (38 samples), endothelial cells (51), embryonic stem cells (51), fibroblasts (46), heart (59) hematopoietic stem/progenitor cells (49), intestine/colon (85), kidney (57), liver (107), lung (44), macrophage/monocyte (170), brain (90), skeletal muscle (118), and T cells (38). FASTQ files were preprocessed into gene x sample expression matrices as described in the above pipeline.

For each query tissue type, a random forest classifier was trained on 2/3 of the training samples from each cell/tissue type using an adapted top-scoring pairs algorithm as described(Peng et al., 2020; Tan and Cahan, 2019). Briefly, template matching was used to identify the top 100 genes highly correlating with each cell/tissue type. These genes were used as input to a gene pair transformation, and each gene pair was ranked by its ability to discriminate between cell/tissue types. Using the top 100 discriminative gene pairs, a gene pair x sample binary matrix was constructed. The random “garbage collection” cell type was generated by randomly shuffling values in the binary matrix, randomly sampling 70 profiles, and appending these to the binary matrix. This matrix was used to train a random forest classifier of 2000 trees, with stratified sampling of 25 samples from each cell type to ensure balanced training among cell/tissue types.

For each cell/tissue type, validation of the classifier was performed on the remaining 1/3 of the training samples and on 60 randomly shuffled expression profiles. Classifier was assessed using area under the precision-recall curve.

### Differential expression and gene set enrichment analysis

Differential expression analysis for RNA-seq data was performed with DESeq2 (Love et al., 2014). Gene set enrichment analysis was performed with Fast GSEA (FGSEA) (Korotkevich et al., 2021), using the C2 and C5 gene sets from the Molecular Signatures Database (MSigDB) (Liberzon et al., 2011; Subramanian et al., 2005).

### Transcriptional regulator scoring

GRN construction was performed as previously described (Radley et al., 2017), and for each cell/tissue type the GRN was subset based on genes existing in all corresponding query samples. For each sample, candidate transcriptional regulators for improving the classification of its target cell/tissue type were scored and prioritized as previously described (Peng et al., 2020):

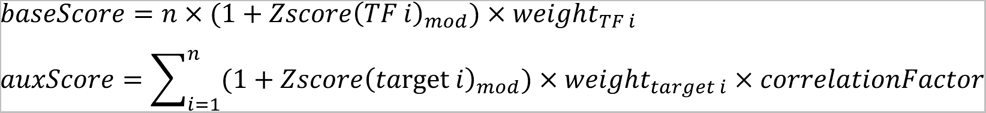

where:

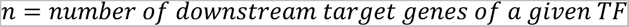

The base score of each TF was determined based on a modified z-score and weight for the TF, and the auxiliary score for the TF’s targets was determined based on a modified z-score, weight, and correlationFactor for each target, where:

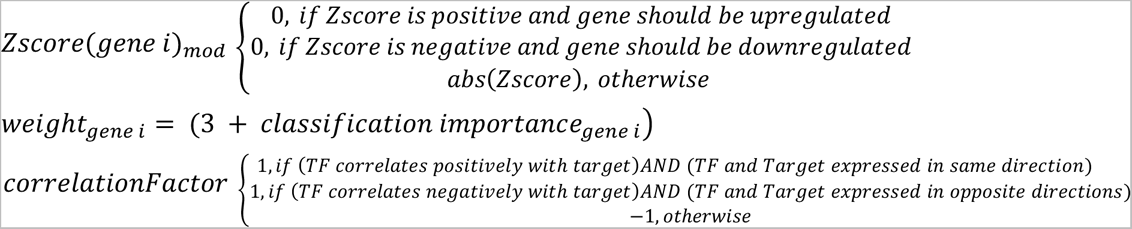

The modified z-score prioritizes genes whose expression patterns do not match those of the primary training samples. In order to consider target genes in order of influence, the weight of each target gene depends on its importance to random forest classifier. Lastly, the correlation factor penalizes cases where targeting a TF would cause incorrect directionality of expression of target genes in the subnetwork.

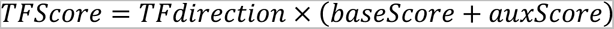

The aggregate TFScore was calculated based on the baseScore and auxScore for a TF, as well as the proposed directionality of the TF to improve classification.

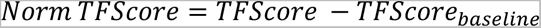

where TFScore_baseline_ is calculated in the same fashion as the TFScore with the modified z-scores for TF and target genes set to 0:

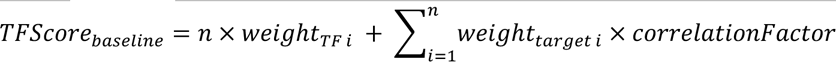

The aggregate TFScore was finally normalized to the TFScore_baseline_ to account for the variation in number of downstream target genes, which would artificially affect the raw TFScore.

### Derivation of primary liver organoids

Primary liver organoids were derived as previously described (Broutier et al., 2016; Huch et al., 2015). Donor liver tissue sample was received directly after surgical procedure on ice in a 50mL tube submerged in PBS -/-. Tissue was immediately transferred to a 10cm petri dish in cold basal medium (Advanced DMEM/F-12 containing 100U/mL penicillin, 100ug/mL streptomycin, 2mM Glutamax, and 10mM HEPES) and kept on a cool surface under a biosafety cabinet. A sterile scalpel was used to cut 0.5-1cm^3 cubes of tissue. Two pieces of tissue were separated and further minced very finely with the scalpel to make cells accessible for digestion. Tissue was transferred to 15mL tube and washed twice with wash medium (DMEM containing 1% FBS, 100U/mL penicillin, 100ug/mL streptomycin, 2mM Glutamax), with supernatant being manually aspirated with a serological pipette after allowing the minced tissue to settle. The tissue was digested with the addition of 4mL digestion solution (HBSS+/+, 1.25 mg/mL collagenase IV, 0.1 mg/mL DNaseI) on a rotor at 37C for 30min. Digested cells suspension was brough to 15mL with cold wash medium and passed through a 70um mesh into a 50mL tube. Cells were pelleted at 300g for 5 min at 8C, the supernatant gently aspirated, and the cells resuspended in another 15mL of cold wash medium, transferred to a 15mL tube and spun again at 300g for 5 min at 8C. The cells were washed an additional two times in cold wash medium and a third time in cold basal medium. The medium was aspirated gently with a serological pipette and a fraction of the cells were transferred to cold growth factor reduced Matrigel at a volume ratio of 1:3, aiming for a concentration of ∼40k cells/mL. Droplets of 25uL volume were distributed into the centers of 48-well-plate wells. The plates were flipped upside down and placed in the tissue culture incubator (37 °C, 5% CO2) for 10-20min until the Matrigel completely solidified. Cultures were overlaid with isolation medium (basal medium supplemented with 1:50 B27 supplement without vitamin A, 1:100 N2 supplement, 1.25 mM N-acetylcysteine, 10% (vol/vol) RSPO1-conditioned medium, 10 mM nicotinamide, 10 nM recombinant human [Leu]-gastrin I, 50 ng/ml recombinant human EGF, 100 ng/ml recombinant human FGF10, 25 ng/ml recombinant human HGF, 10 μM Forskolin and 5 μM A83-01, 25 ng/ml recombinant human Noggin, 30% v/v Wnt3a-conditioned medium and 10 μM Y-27632) for 4 days before switching to expansion medium (same as isolation medium but without Noggin, Wnt3A-conditioned medium, and Y-27632), and changing medium every 3-4 days. Cultures were passaged on day 13 after isolation.

### Passaging of human liver organoids

Organoids were passaged 13 days after the original isolation and every 5-7 days afterwards. Matrigel was thawed ahead of time and kept on ice. Organoid cultures were disrupted with a P1000 micropipette and the wells rinsed with twice with 500uL of cold basal medium to dissolve the Matrigel droplet and transfer the cells to a 15mL centrifuge tube. The tubes were topped off to 13mL with cold basal medium and pipetted up and down 5 times to completely dissolve the Matrigel. Organoids were spun at 200g for 5 min at 8C and aspirated until there was 1mL remaining. The organoids were pipetted repeatedly using a P1000 micropipette until the organoids were sufficiently dissociated. The tubes were again topped to 13mL with basal medium, spun at 200g for 5 min at 8C, and aspirated. The cells were mixed with the Matrigel to reach a passage ratio of 1:6 and droplets were formed and overlaid as with the isolation procedure, but with expansion medium instead of isolation medium.

### Immunofluorescence staining and imaging

Whole mount staining of primary human liver organoids was performed as follows. Glass bottom plates (Ibidi) containing Matrigel droplet organoid suspensions were rinsed twice with cold PBS -/- (Quality Biological), then fixed in 4% paraformaldehyde (Electron Microscopy Sciences) for 30 minutes at room temperature (RT); remaining incubations and washes were performed at RT unless otherwise specified. Wells were rinsed 3x with PBS -/-, then permeabilized with 0.3% TritonX-100 for 25 minutes. Wells were again rinsed 3x in PBS -/- and blocked with 5% normal donkey serum (Jackson Immunolabs) for 45 minutes, rinsed 1x in PBS -/- and incubated overnight at 4C with primary antibodies (see supplementary table for antibody list and dilutions) in 5% donkey serum in 0.05% Tween-20 in PBS-/- (PBST). A negative control was incubated under the same conditions but with no primary antibodies. Cells were washed with PBST 3x for 10 minutes per wash, then incubated with secondary antibody in PBST for 90 minutes. Wells were rinsed 2x with PBS then incubated with a 1:1500 dilution of Hoechst (Invitrogen) in PBST for 10 minutes before a final 3 washes of 10 minutes each. The stained cultures were then imaged on a Nikon A1 spectral confocal using z-stack scans, which were later processed in ImageJ/FIJI to generate z-projections shown in the figures. The negative control that was incubated with secondary antibodies, but not primary antibodies, was used to set the microscope parameters to determine the background threshold for each channel. Antibody catalog numbers and dilutions can be found in Table S12.

## Acknowledgments

E.K.L. is supported by the National Cancer Institute of the National Institutes of Health under award F31CA250489 and by the ARCS Foundation, Metro-Washington Chapter through the JCM Foundation. M.R.E. is supported by R01s from the National Institute of Biomedical Imaging and Bioengineering (EB028532) and the National Heart Lung and Blood Institute (HL141805), National Science Foundation, CBET, Award Number 2134999 and the Pittsburgh Liver Research Center (P30DK120531). PC is supported by EB028532 and R35GM124725. The graphical abstract was created with Biorender.com.

## Author Contributions

E.K.L curated data, wrote code, performed analysis, and wrote the manuscript. J.V. performed experiments and edited the manuscript. D.P. wrote code and edited the manuscript. C.K guided analysis and edited the manuscript. M.E. oversaw experiments, guided analysis, and edited the manuscript. P.C. conceived and oversaw the project, guided analysis, and edited the manuscript.

---

**Figure S1.**
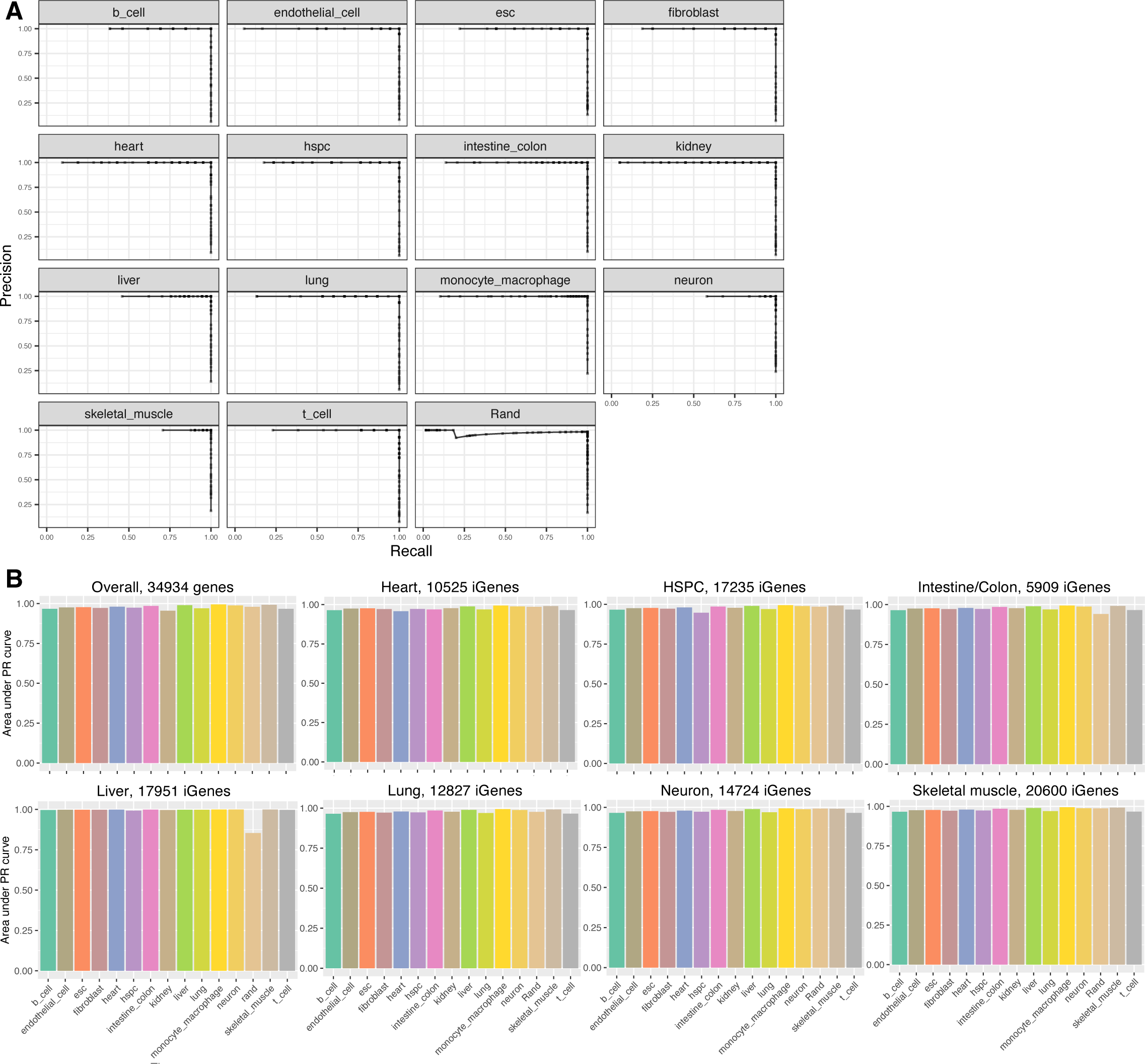
Classifier performance by precision and recall. Related to Figure 1. (A) Precision-recall (PR) curves for each cell/tissue type-specific classifier. For each curve, each point represents a classification score threshold for a sample to be classified as the given cell/tissue type. High precision indicates low false positive rate, and high recall indicates low false negative rate. (B) Area under the PR curve (AUPR) for classifiers trained on different subsets of intersecting genes (iGenes). For example, the plot on the first row, second column indicates the AUPR for classifiers trained on 10525 iGenes included across all publicly available heart engineering studies.

**Figure S2.**
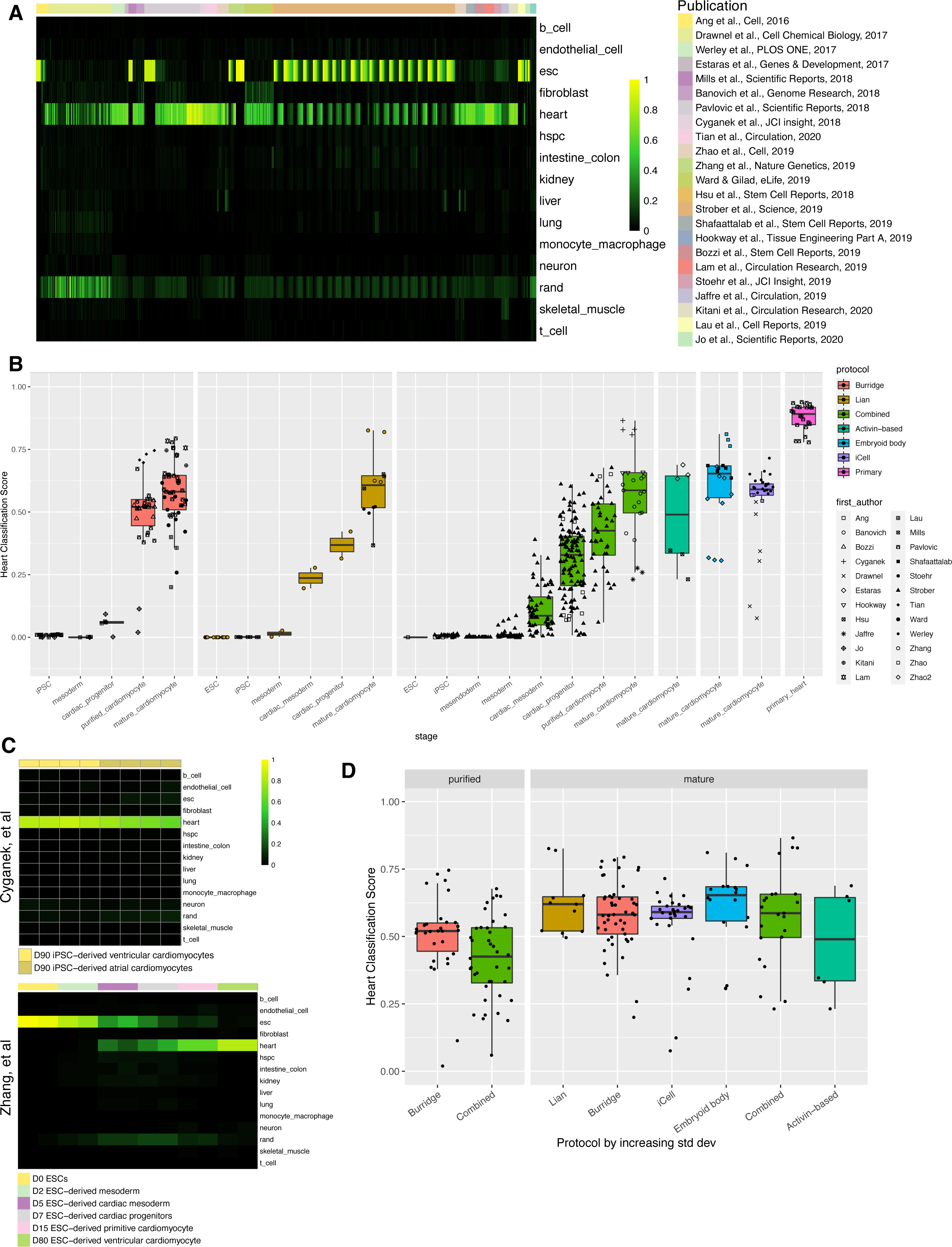
Cross-study meta-analysis of cardiomyocyte engineering protocols. Related to Figure 2. (A) Compiled classification heatmap of samples from curated publicly available cardiomyocyte engineering studies. (B) Heart classification scores for each sample stratified by cardiomyocyte engineering protocol. Combined: Combination of Lian protocol with additional metabolic selection step. (C) Heatmap of classification scores for the two studies, Cyganek et al (Combined Protocol) and Zhang et al (Lian Protocol + MYL2 sorting), which achieved the most highly classifying cardiomyocytes. (D) Heart classification scores for purified (metabolically selected) or “mature” derived cardiomyocytes only, grouped by protocol. For (B) and (D), samples were limited to healthy/unperturbed purified and mature CMs (from the the top performing protocol variant, if relevant) per study. Within each facet, protocols are ordered by increasing standard deviation of classification score.

**Figure S3.**
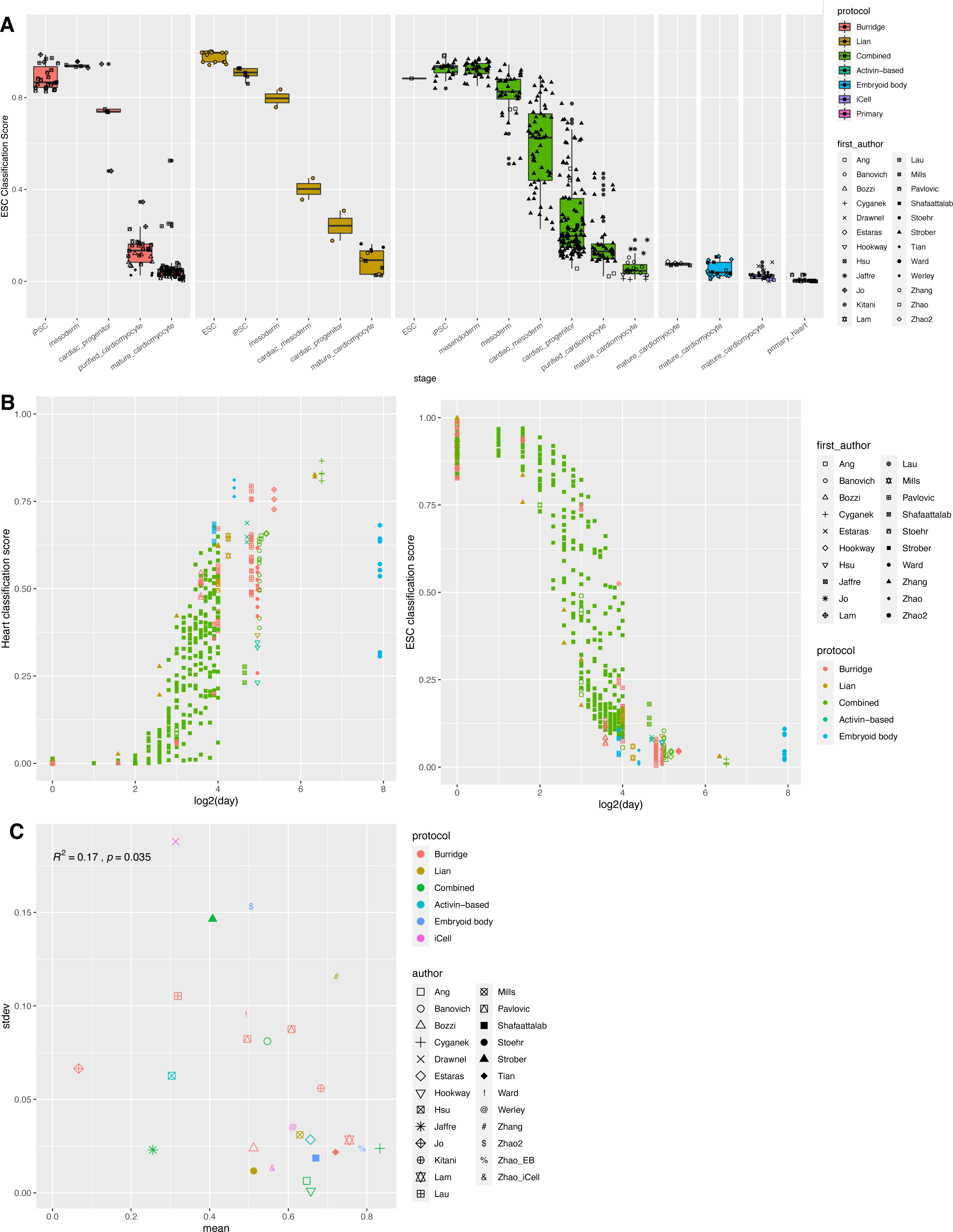
Global patterns of ESC signature in cardiomyocyte engineering protocols. Related to Figure 2. (C) Embryonic stem cell (ESC) classification scores for each sample stratified by cardiomyocyte engineering protocol. (D) Heart classification vs day (log scale, plotted only for samples/studies that provided time points) (left). ESC classification vs day (log scale, plotted only for samples/studies that provided time points) (right). (E) Correlation of classification mean and standard deviation for each study. For (B) and (C), samples were limited to healthy/unperturbed purified and “mature” CMs (from the the top performing protocol variant, if relevant) per study.

**Figure S4.**
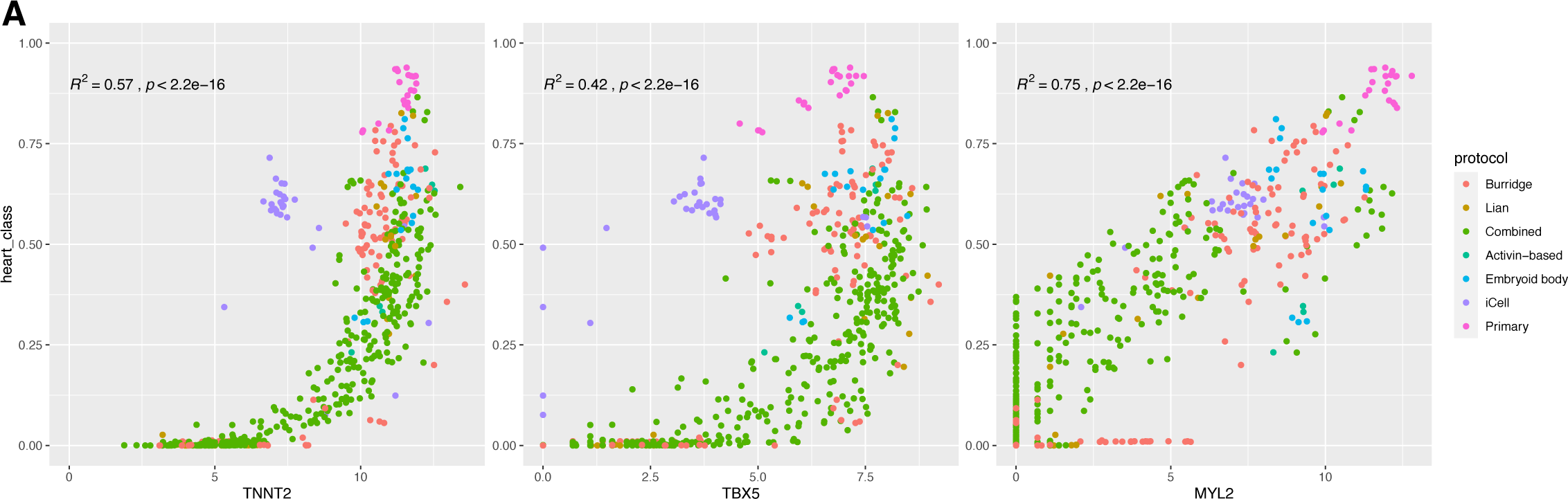
Correlation of heart classification with canonical heart marker genes. Related to Figure 2. (A) Scatterplot of heart classification score vs cardiomyocyte marker gene (TNNT2, TBX5, and MYL2) expression (log-transformed counts) for each engineered (or primary control) cardiomyocyte sample, grouped by protocol.

**Figure S5.**
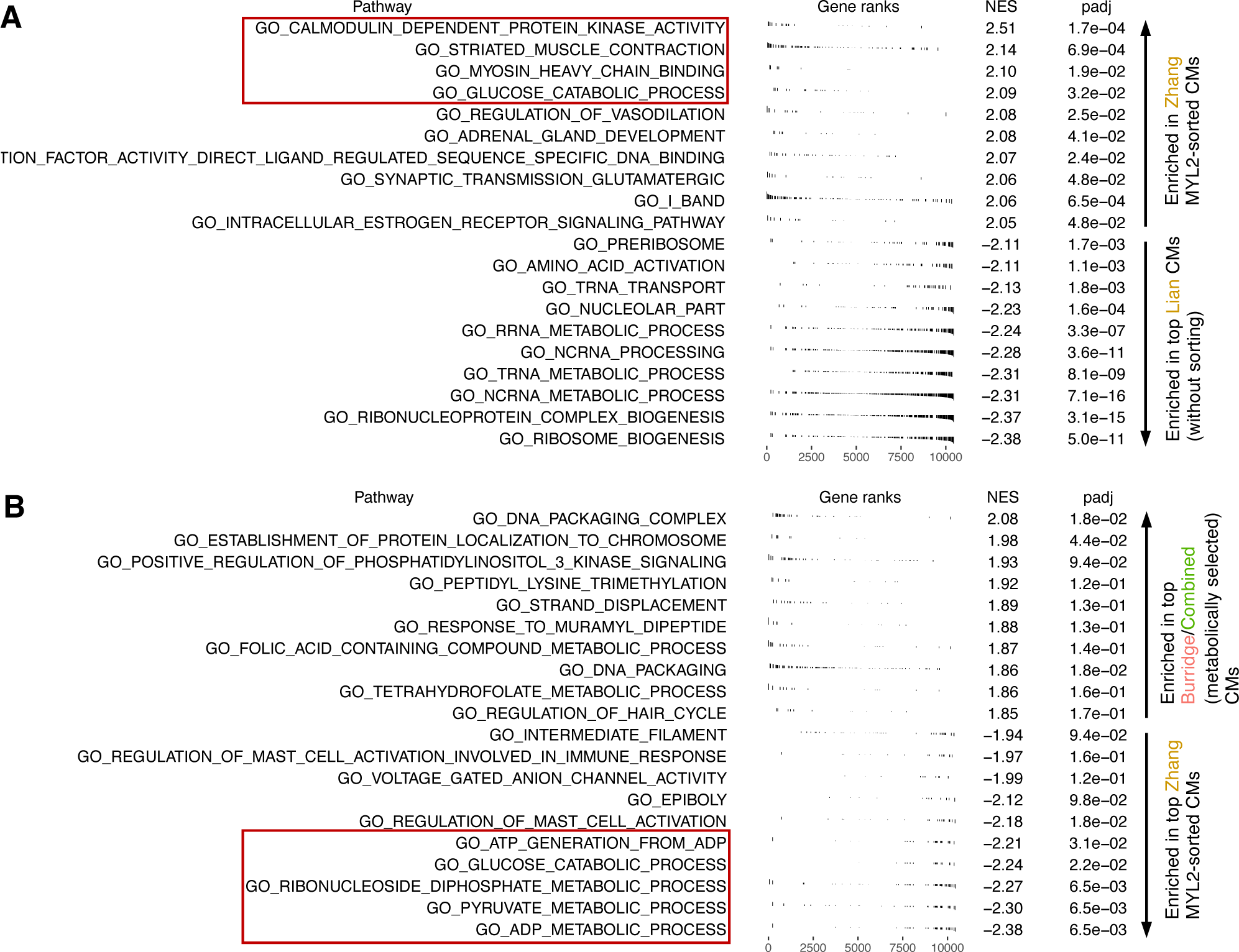
Gene expression patterns in top-performing studies. Related to Figure 2. (A) Summary plot for GSEA performed on differentially expressed genes between Zhang MYL2-sorted CMs (positive NES) vs top-classifying Lian CMs (Zhang unsorted samples, Mills, Stoehr; negative NES) without sorting. (B) Summary plot for GSEA performed on differentially expressed genes between top-classifying CMs derived via Burridge and Combined protocols (positive NES) vs Zhang MYL2-sorted CMs (negative NES). NES: normalized enrichment score. padj: Adjusted p-value based on Benjamini-Hochberg correction.

**Figure S6.**
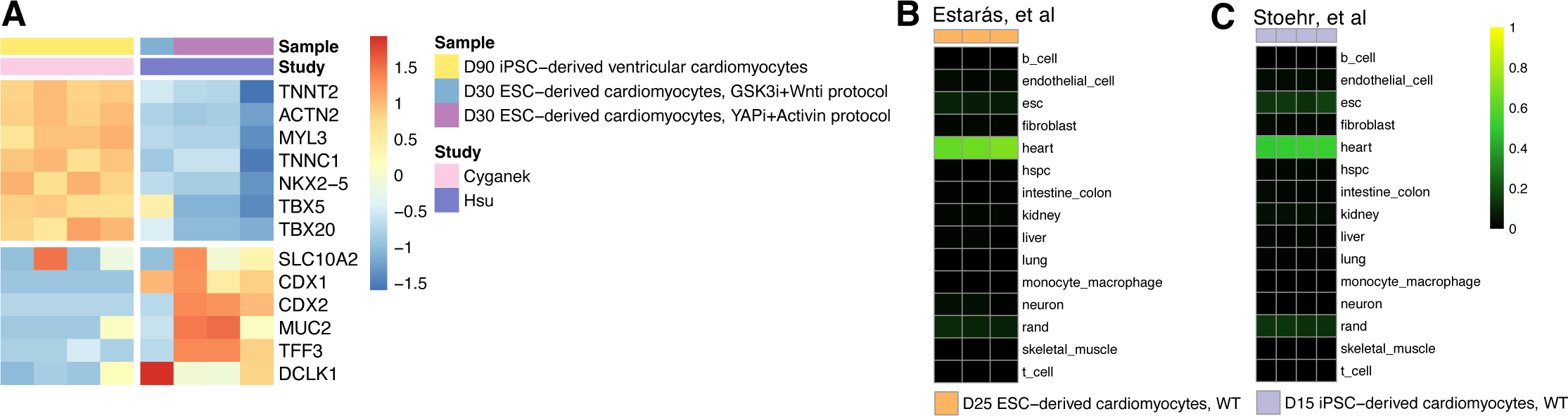
Off-target signatures in cardiomyocyte studies. Related to Figure 3. (A) Heatmap of gene expression for canonical cardiomyocyte and intestinal marker genes for the study with the most highly classifying cardiomyocytes (Cyganek et al) compared to Hsu et al study. Heatmap color scale: per gene z-score of log transformed TPM. (B) Heatmap of classification scores for Estaras et al study, who used the one-step activin differentiation protocol followed by metabolic selection (C) Heatmap of classification scores for Stoehr et al study.

**Figure S7.**
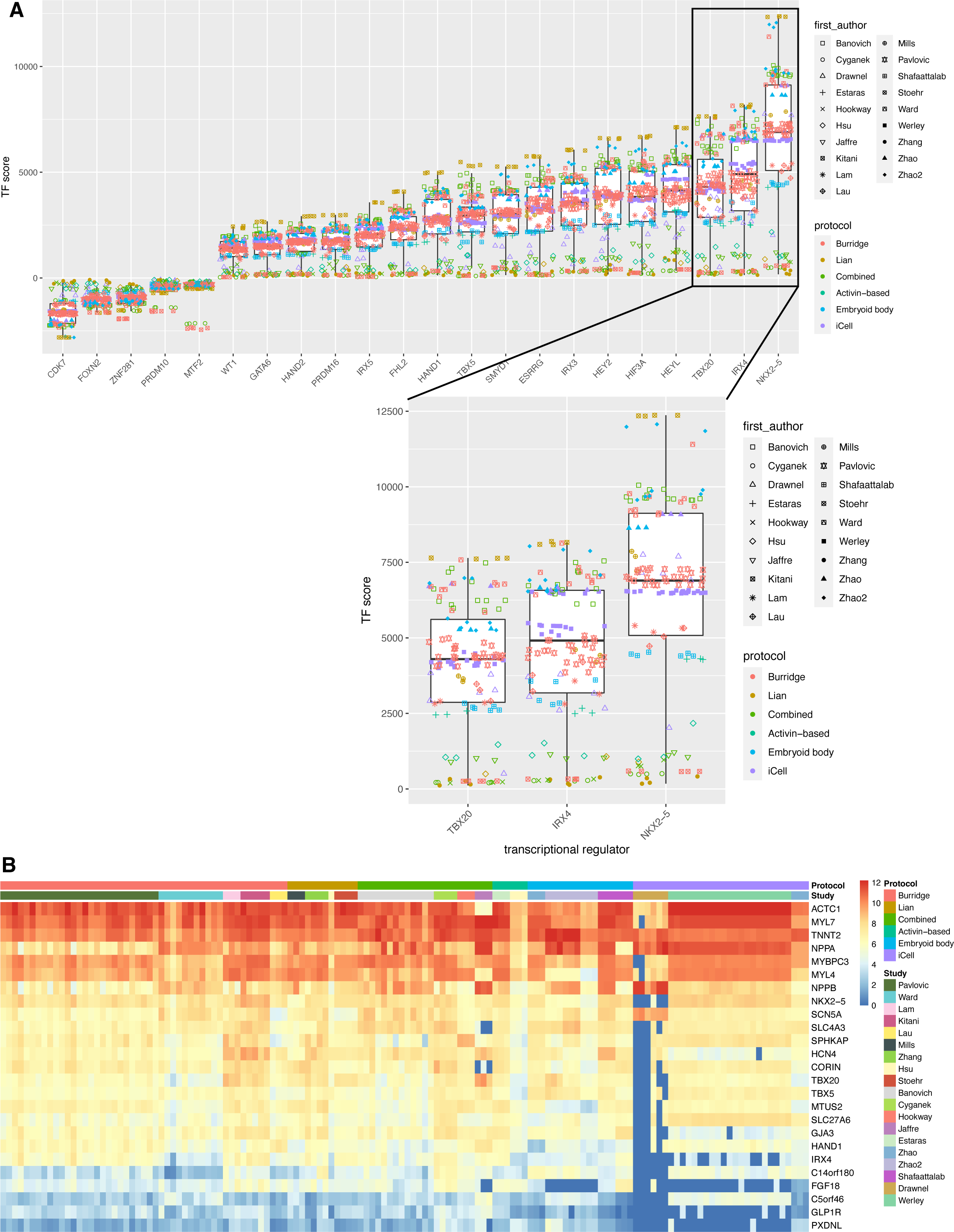
Scoring heart transcriptional regulators for protocol improvement. Related to Figure 2. (A) Compiled boxplots of the top 25 (by greatest mean magnitude) heart transcriptional regulator scores of mature engineered samples as calculated by the NIS algorithm. A positive value indicates the need for upregulation in a given sample; a negative value indicates the need for downregulation. Inset: TF scores for the top 3 implicated transcriptional regulators. Samples were limited to healthy/unperturbed purified and mature CMs (from the the top performing protocol variant, if relevant) per study. (B) Quantile-normalized counts for predicted targets of NKX2-5 in mature engineered cardiomyocytes, ranked by mean gene expression across samples.

**Figure S8.**
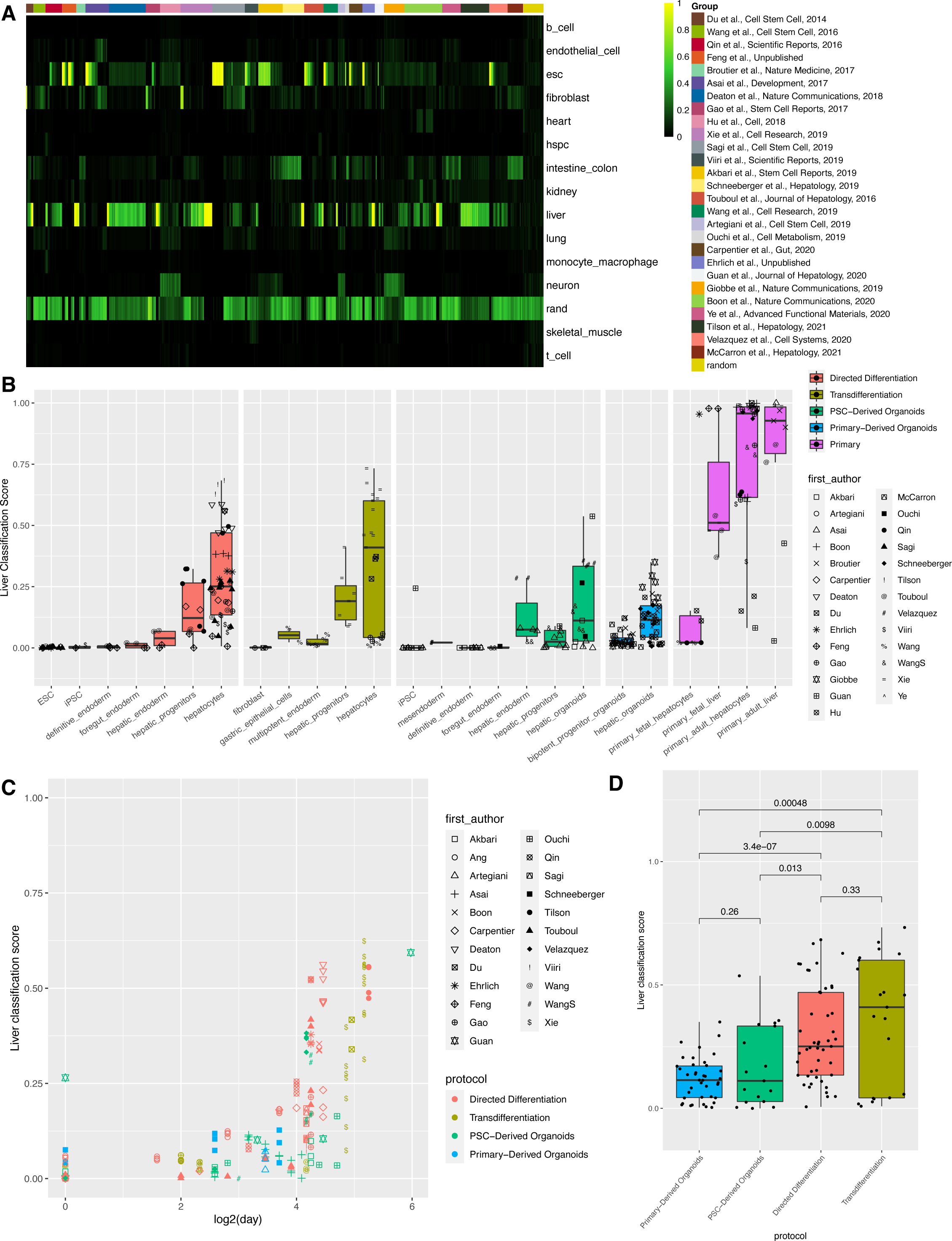
Cross-study meta-analysis of hepatocyte engineering protocols. Related to Figure 4. (A) Compiled classification heatmap of samples from curated publicly available hepatocyte engineering studies. (B) Liver classification scores for each sample stratified by hepatocyte engineering protocol. (C) Liver classification score versus day (log scale) for engineered samples where a time point was provided. (D) Liver classification scores for mature derived hepatocytes and hepatic organoids only, grouped by protocol and ordered by increasing standard deviation. For (B)-(D), samples were limited to healthy/unperturbed mature or differentiated hepatocytes and hepatic organoids (from the the top performing protocol variant, if relevant) per study.

**Figure S9.**
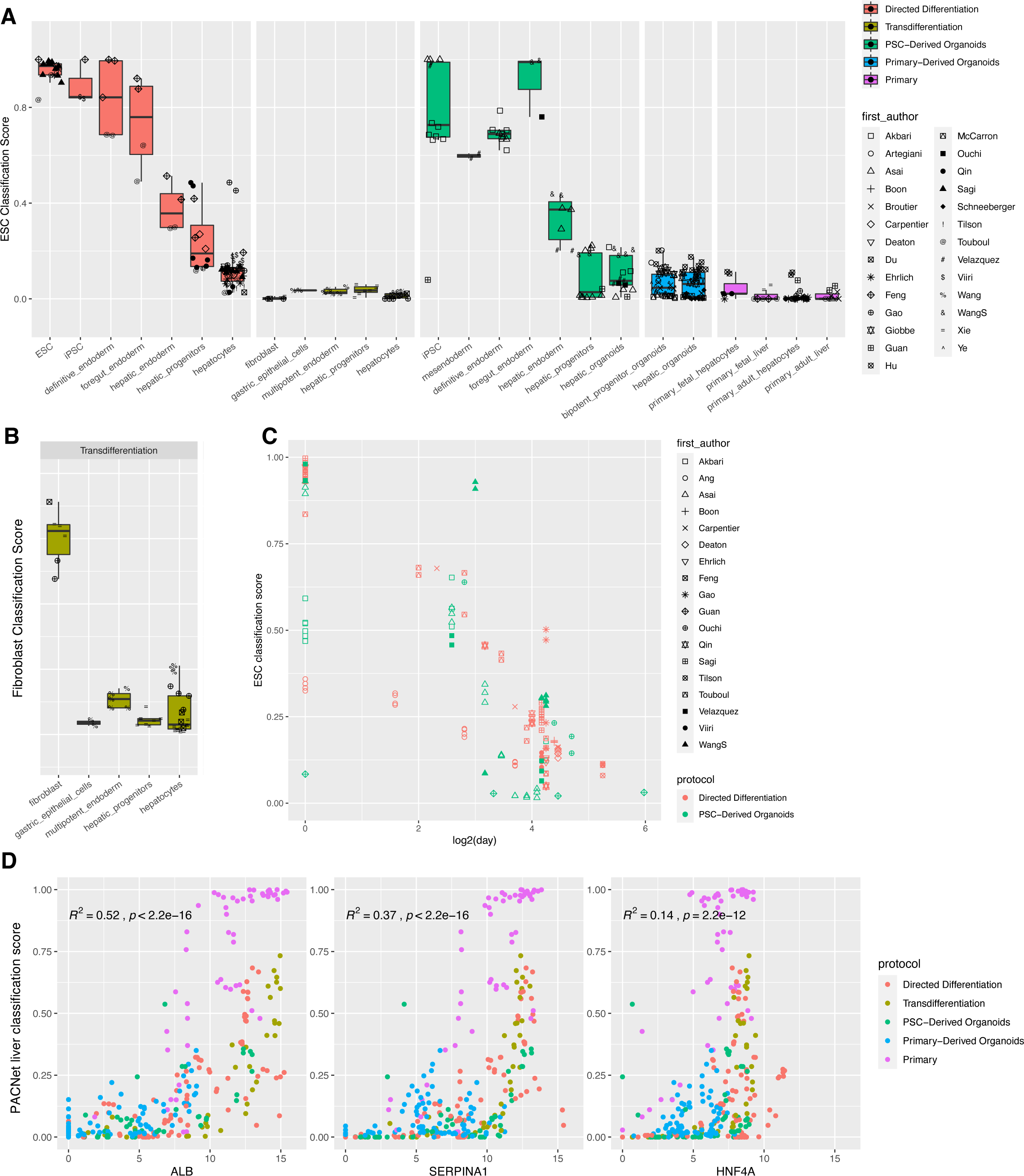
Global performance of cardiomyocyte engineering protocols assessed via classification. Related to Figure 4. (A) ESC classification scores for each sample stratified by hepatocyte engineering protocol. (B) Fibroblast classification score included for studies in the transdifferentiation category (bottom right). (C) ESC classification score versus day (log scale) for engineered samples for which a time point was provided, starting from the ESC or iPSC stage. (D) Scatterplot of liver classification score vs hepatocyte marker gene expression (ALB, SERPINA1, HNF4A, log-transformed counts) for each engineered hepatocyte sample. For (A) and (B), samples were limited to healthy/unperturbed mature or differentiated hepatocytes and hepatic organoids (from the the top performing protocol variant, if relevant) per study.

**Figure S10.**
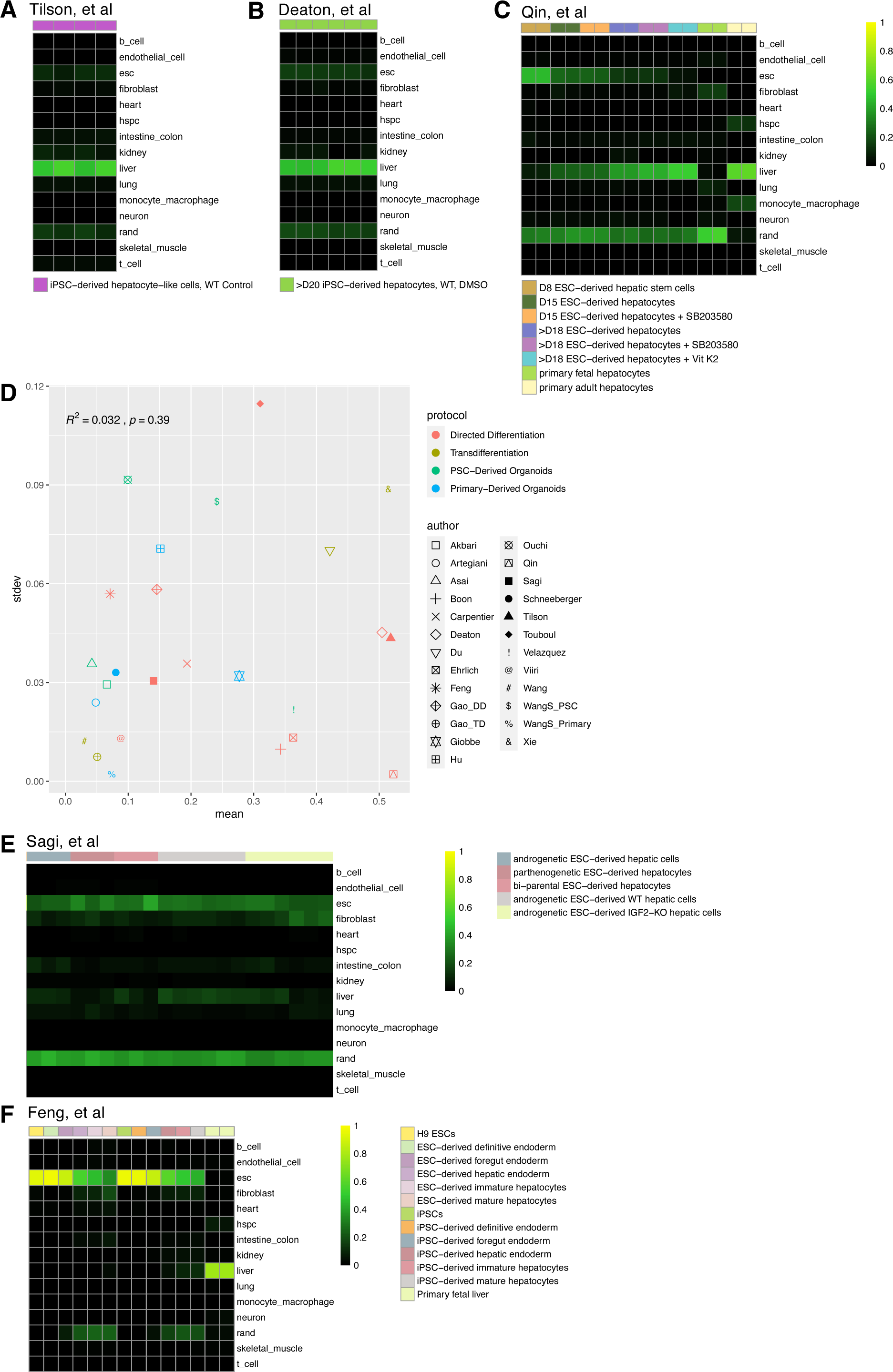
Notable hepatocyte derivation studies and patterns. Related to Figure 4. Heatmap of classification scores for (A) Tilson et al, (B) Deaton et al, and (C) Qin et al, which produced some of the most highly classifying hepatocyte samples. (D) Correlation of classification mean and standard deviation for each study. Samples were limited to healthy/unperturbed mature or differentiated hepatocyte and hepatic organoid samples (from the the top performing protocol variant, if relevant) per study. Heatmap of classification scores for two studies, (E) Sagi et al and (F) Feng et al, which produced hepatocyte samples with a substantial residual ESC signatures.

**Figure S11.**
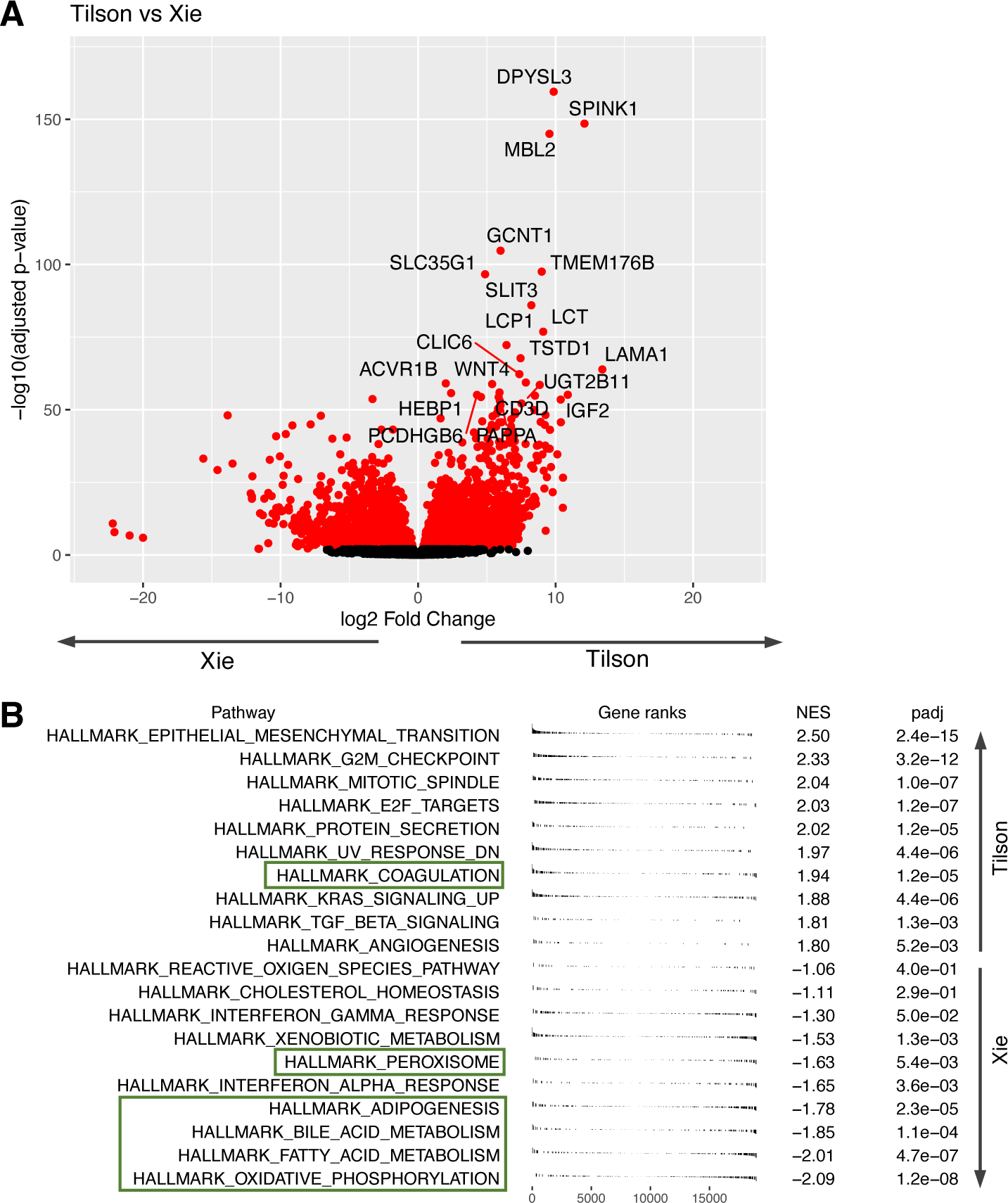
Gene expression differences between the top-performing hepatocyte derivation studies. Related to Figure 4. (A) Volcano plot of differentially expressed genes comparing hepatocytes derived by Tilson (positive log2 fold change) vs by Xie (negative log2 fold change). Red: adjusted p-value < 0.05. (B) Summary plot of GSEA comparing hepatocytes derived by Tilson (positive NES) vs by Xie (negative NES). Green boxes denote liver-related gene sets of interest. NES: normalized enrichment score. padj: Adjusted p-value based on Benjamini-Hochberg correction.

**Figure S12.**
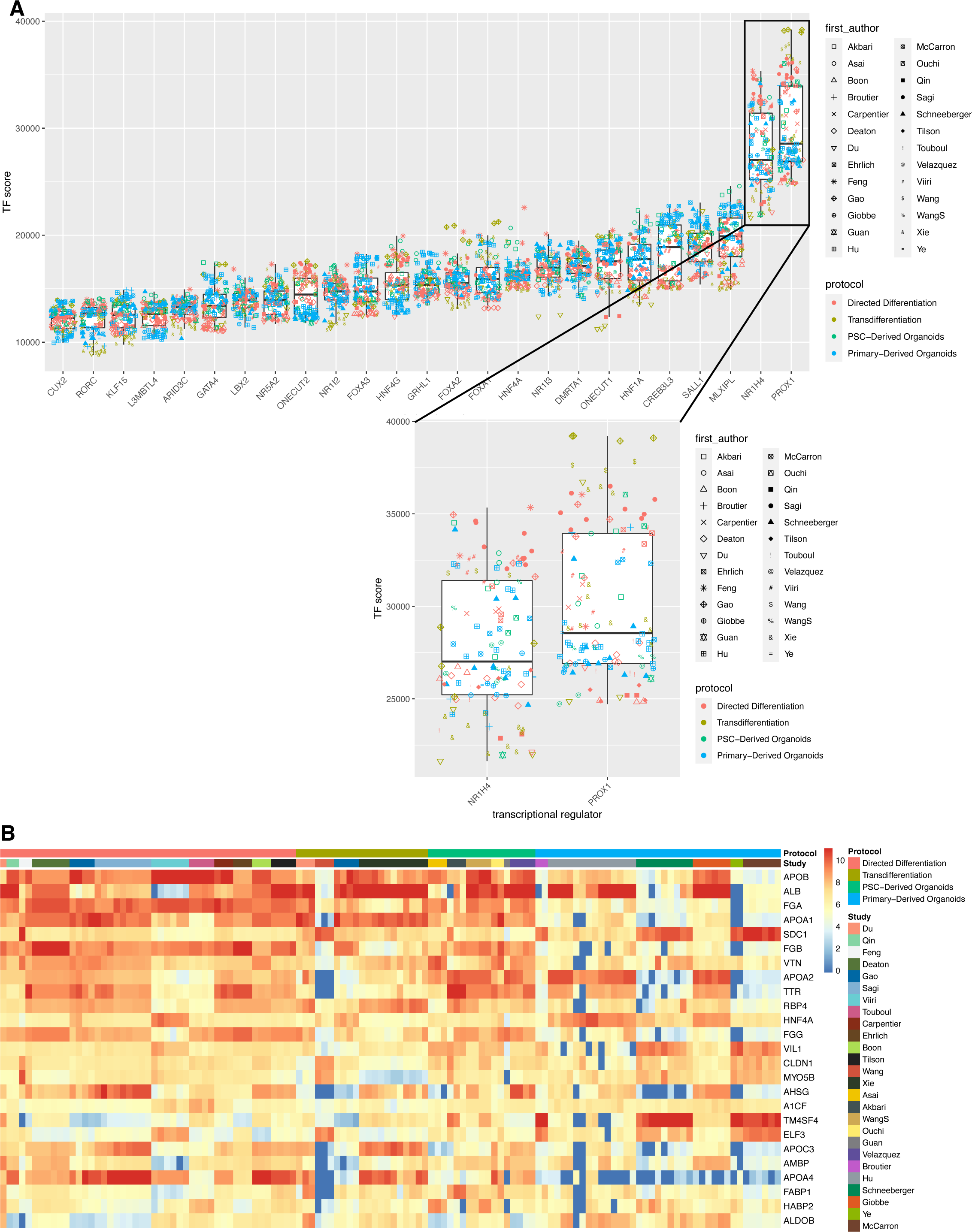
Scoring liver transcriptional regulators for protocol improvement. Related to Figure 4. (A) Compiled boxplots of the top 25 (by greatest mean magnitude) liver transcriptional regulator scores of mature engineered samples as determined by the NIS algorithm. A positive value indicates the need for upregulation in a given sample; a negative value indicates the need for downregulation. Inset: TF scores for the top 2 implicated transcriptional regulators. Samples were limited to healthy/unperturbed mature or differentiated hepatocytes and hepatic organoids (from the the top performing protocol variant, if relevant) per study. (B) Quantile-normalized counts for predicted targets of NR1H4 in mature engineered hepatocytes and hepatic organoids only, ranked by mean gene expression across all samples.

**Figure S13.**
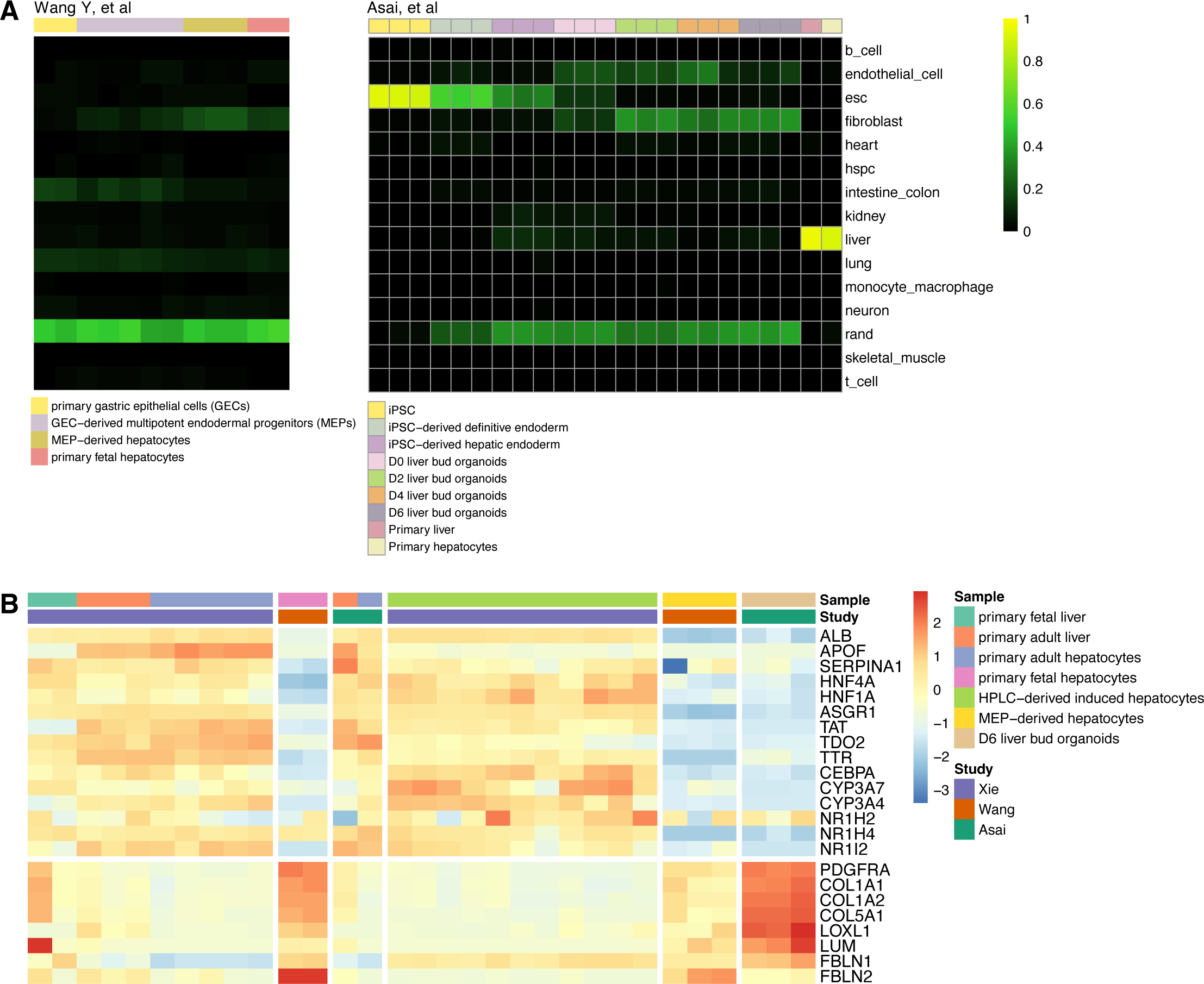
Off-target fibroblast signatures in engineered hepatic cells. Related to Figure 5. (A) Classification heatmaps for hepatocyte derivation studies performed by Wang et al (left) and Asai et al (right). (B) Heatmap of hepatocyte and fibroblast marker gene expression for highest classifying hepatocyte derivation study (Xie et al) vs Wang et al and Asai et al. Heatmap color scale: per gene z-score of log-transformed TPM.

**Figure S14.**
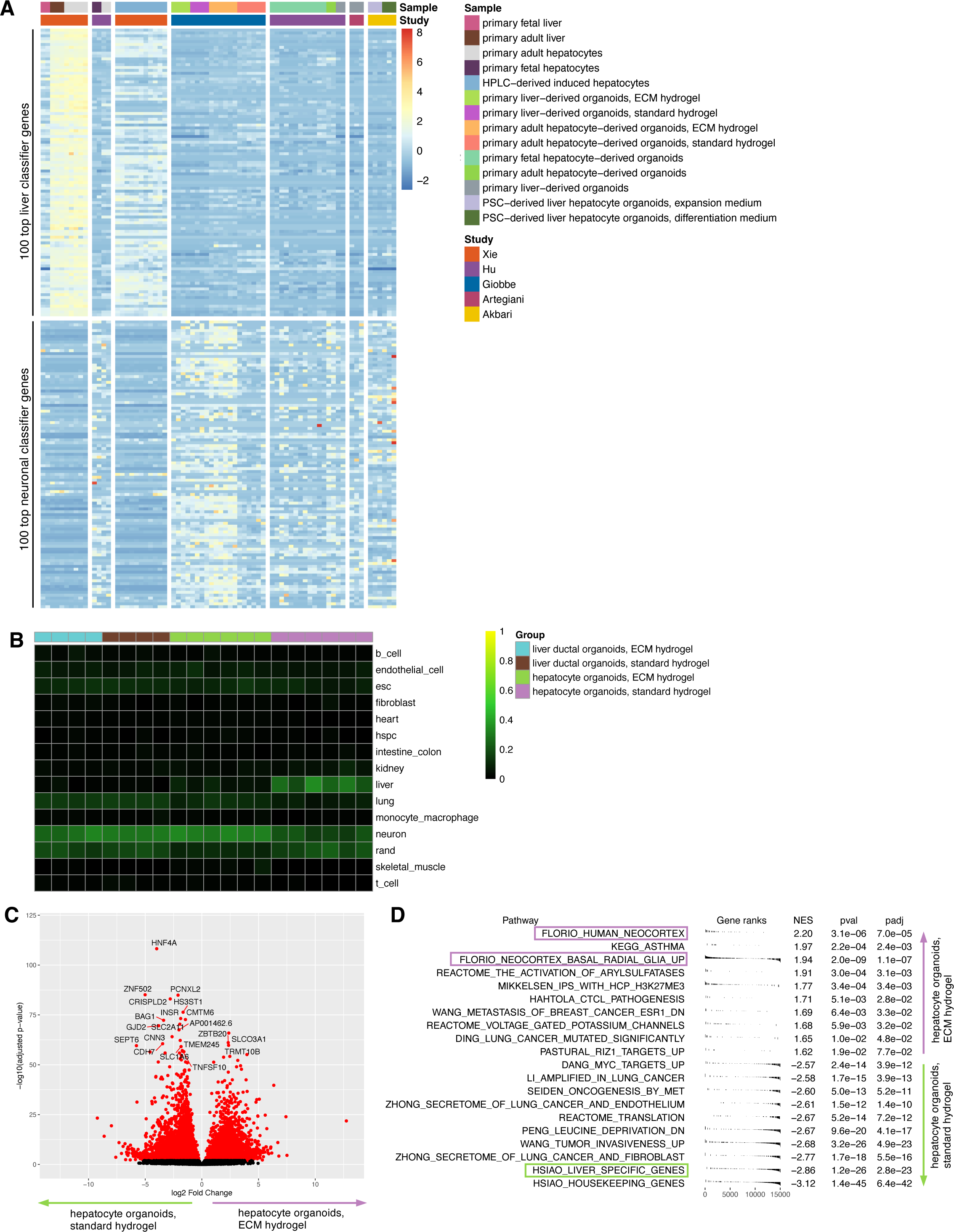
Off-target neuronal signatures in primary hepatocyte-derived organoids. Related to Figure 6. (A) Expression heatmap of the 100 top genes contributing to the liver (top) and neuronal (bottom) random forest classifiers. Heatmap color scale: per gene z-score of log-transformed TPM. (B) Classification heatmap for primary liver-derived organoids generated by Giobbe et al. (C) Volcano plot of differentially expressed genes comparing hepatocyte organoids in ECM hydrogel (positive log2 fold change) and hepatocyte organoids in standard hydrogel (negative log2 fold change) generated by Giobbe et al. Red: adjusted p-value < 0.05. (D) Summary plot of GSEA comparing hepatocyte organoids in ECM hydrogel (positive NES) and hepatocyte organoids in standard hydrogel (negative NES). Boxes denote neural and liver gene sets of interest. NES: normalized enrichment score. padj: Adjusted p-value based on Benjamini-Hochberg correction.

**Figure S15.**
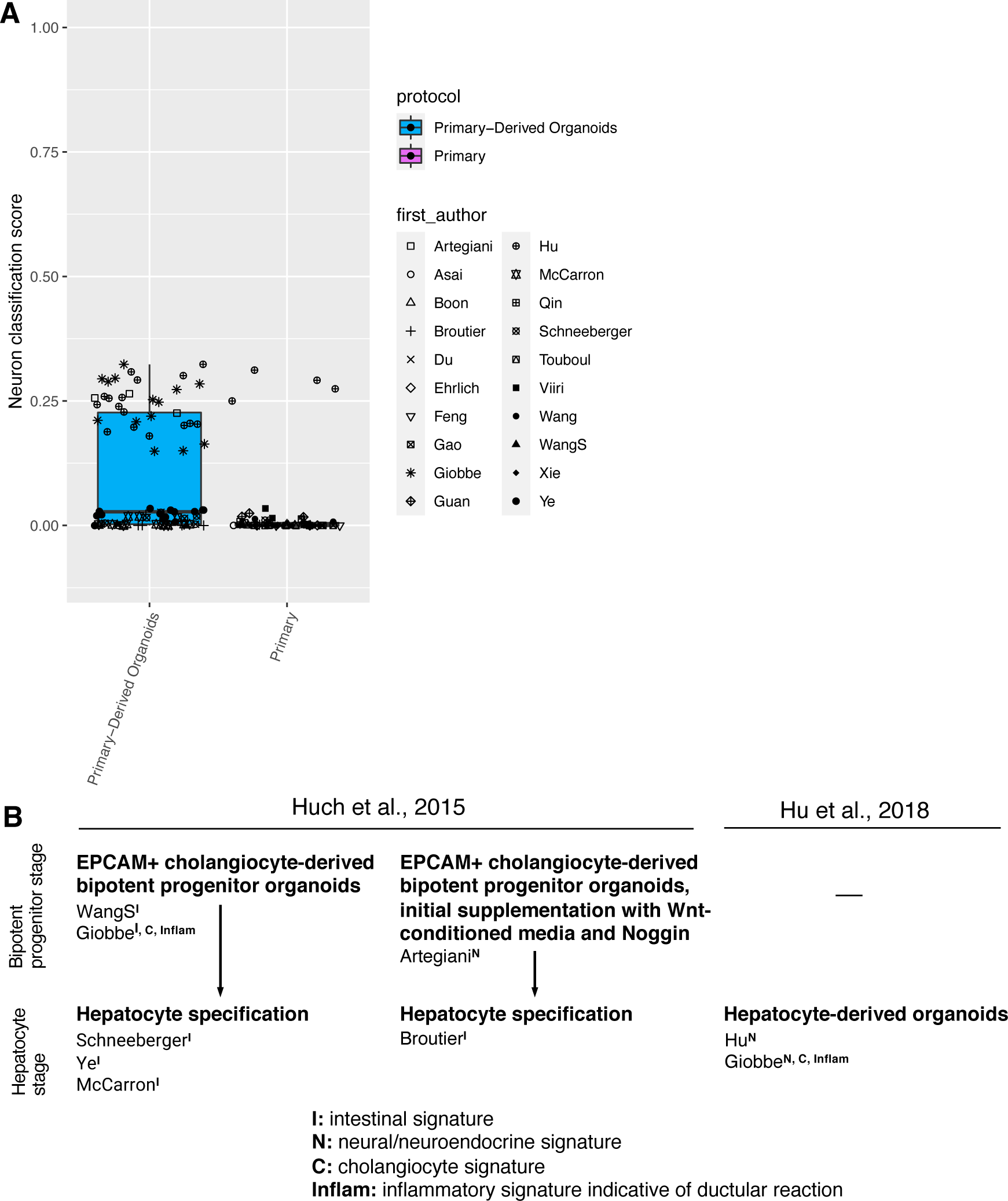
Patterns in off-target signatures in primary hepatocyte-derived organoids. Related to Figure 6. (A) Neuronal classification scores for primary-derived organoid samples vs primary liver or primary hepatocyte control samples. (B) Schematic of differences in derivation strategies for primary liver-derived organoids denoting detected off-target signatures.

**Figure S16.**
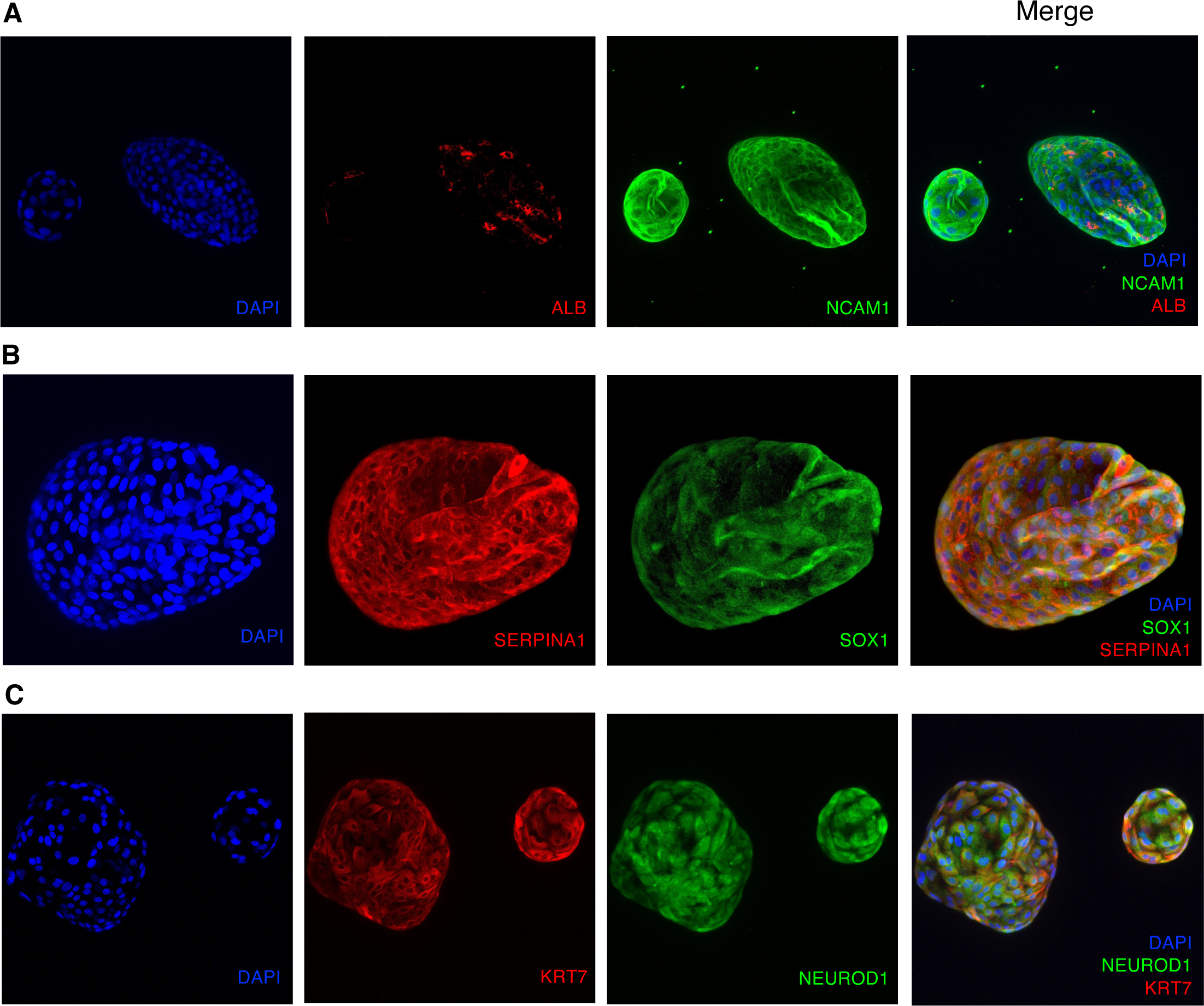
Immunofluorescence staining for off-target neuronal signatures in primary hepatocyte-derived organoids. Related to Figure 6. Immunofluorescence staining, single channel and merge, for (A) hepatocyte marker Albumin (ALB) and neuroendocrine marker NCAM1; (B) hepatocyte marker SERPINA1 and neuronal marker SOX1; and (C) cholangiocyte marker KRT7 and neuronal marker NEUROD1. Blue: DAPI.

## Supplemental Tables

**Table S1. PACNet human training studies and samples for 14 tissue types. Related to Figure 1.**

**Table S2. PACNet human engineered reference studies for heart, hematopoietic stem and progenitor cells, intestine/colon, liver, lung, neuron, and skeletal muscle. Related to Figure 1.**

**Table S3. Benchmarking PACNet classification scores using alternate RNAseq preprocessing methods. Related to Figure 1.**

**Table S4. Panel of engineered cardiomyocyte samples. Related to Figures 2 and 3.**

**Table S5. Panel of engineered liver samples. Related to Figures 4-6.**

**Tables S6-S10. Panels of engineered samples for HSPCs, intestine/colon, lung, neuron, and skeletal muscle. Related to Figure 1.**

**Table S11. Top 100 genes contributing to the liver and neuronal PACNet classifiers. Related to Figure 6.**

**Table S12. Antibody catalog numbers and dilutions. Related to Figure 6.**

